# Rational design of synthetic proteins using a genome-scale CRISPR screen

**DOI:** 10.64898/2026.02.19.706875

**Authors:** Wells H. Burrell, Simon J. Mueller, Zharko Daniloski, P. Duffy Doyle, Anne B. Rovsing, Christopher James, Max Drabkin, Chien-Yu Chou, Hei Yu Annika So, Lyla Katgara, Akash Sookdeo, Lu Lu, Georges-Ibrahim Cisse, Rachel E. Yan, Neville E. Sanjana

## Abstract

Protein structure prediction using deep learning has revolutionized protein design. Yet, our understanding of protein function remains a key limitation for designing novel proteins that perform complex biological tasks. Here, we adopt a massively-parallel, function-first approach to rationally design synthetic proteins. Using genome-scale CRISPR activation, we overexpress ∼19,000 human proteins and measure their impact on precise gene editing. We identify over 800 native proteins that promote homology-directed repair. Using top candidates, we then design synthetic genome editors — Targeted Repair fUsion Editors (TruEditors) — by fusing full-length proteins or smaller core domains to the Cas9 nuclease. We develop 12 unique TruEditors that improve precise gene editing in diverse cell types and at genomic loci where existing methods for precise gene editing fail. Using affinity proteomics, we show that these synthetic proteins work by coordinating with endogenous DNA repair complexes. The delivery of TruEditors via mRNA more than doubles the rate of chimeric antigen receptor (CAR) insertion into the *TRAC* locus of primary human T cells, enhancing CAR T cell-directed tumor cell killing, and improves precise editing in human pluripotent stem cells more than three-fold. Overall, our study demonstrates that genome-wide protein overexpression screens can guide the rational design of synthetic proteins for specific biological tasks.

## Main

The goal of protein design is to engineer novel proteins with specific, user-defined functions and structures. Recently, several groups have capitalized on deep learning models trained on thousands of protein structures to generate synthetic proteins with specific properties, such as binders for clinically-relevant targets like PD-1 or EGFR^1–6^. Others have paired protein sequences with natural language descriptions and, using large language models, have generated novel sequences with specific functional properties^7^, including programmable CRISPR nucleases similar to the widely-used *S. pyogenes* Cas9^8^.

In contrast, efforts to design gene editors to make precise edits in the human genome have relied primarily on hypothesis-driven fusion of specific nucleic acid-modifying proteins to CRISPR nucleases, resulting in a diverse array of editing modalities^9–14^. All of these editors interact with endogenous DNA repair pathways in order to install precise changes into target genomes^15–17^. Improving upon these tools has hinged on our understanding of the interactions between gene editing and DNA repair proteins^18–29^. In the case of homology-directed repair (HDR), strategies to improve precise gene editing have typically focused on inhibition of the nonhomologous end-joining (NHEJ) pathway, which dominates in mammalian cells^20–23^. This can lead to unintended DNA damage and cytotoxicity due to global inhibition of DNA repair pathways^30^. In contrast, more targeted strategies to locally inhibit NHEJ or promote HDR through the recruitment of DNA repair factors by direct^24–26^ or indirect^27^ fusion to Cas9 can bolster precise editing without global consequences on DNA repair. To date, this has been limited to a handful of previously-characterized DNA-repair proteins^28^.

In this study, we take a function-first approach to design proteins for precise gene editing. Using a massively-parallel CRISPR activation screen, we couple the overexpression of ∼19,000 proteins with a functional readout for precise DNA repair. Using this proteome-scale dataset, we design novel Cas9 fusion proteins with full-length candidates or minimal core domains that locally favor HDR over NHEJ. With these new proteins (TruEditors), we demonstrate improved editing at multiple disease-relevant loci, including edits that are difficult or impossible to make with other precise editing tools like prime or base editing. To understand the molecular mechanisms underlying TruEditors, we map their distinct protein-protein interactions and pinpoint critical residues for protein complex formation with endogenous DNA repair factors. Finally, we demonstrate the utility of several TruEditors to precisely edit human pluripotent stem cells and to engineer primary human T cells for cell-based immunotherapies.

### A proteome-scale functional map for precise DNA editing

To design synthetic proteins for a complex biological task (precise DNA repair), we aimed to systematically identify native proteins from the entire human proteome that enhance Cas9-mediated gene editing outcomes. We began by performing a genome-scale CRISPR activation (CRISPRa) screen with a fluorescent reporter for DNA repair outcomes (**Fig. 1a**). As a functional readout of gene editing, we used a stoplight reporter system where two precise edits in neighboring codons are required to spectrally shift EGFP to EBFP^31^.

**Figure 1.**
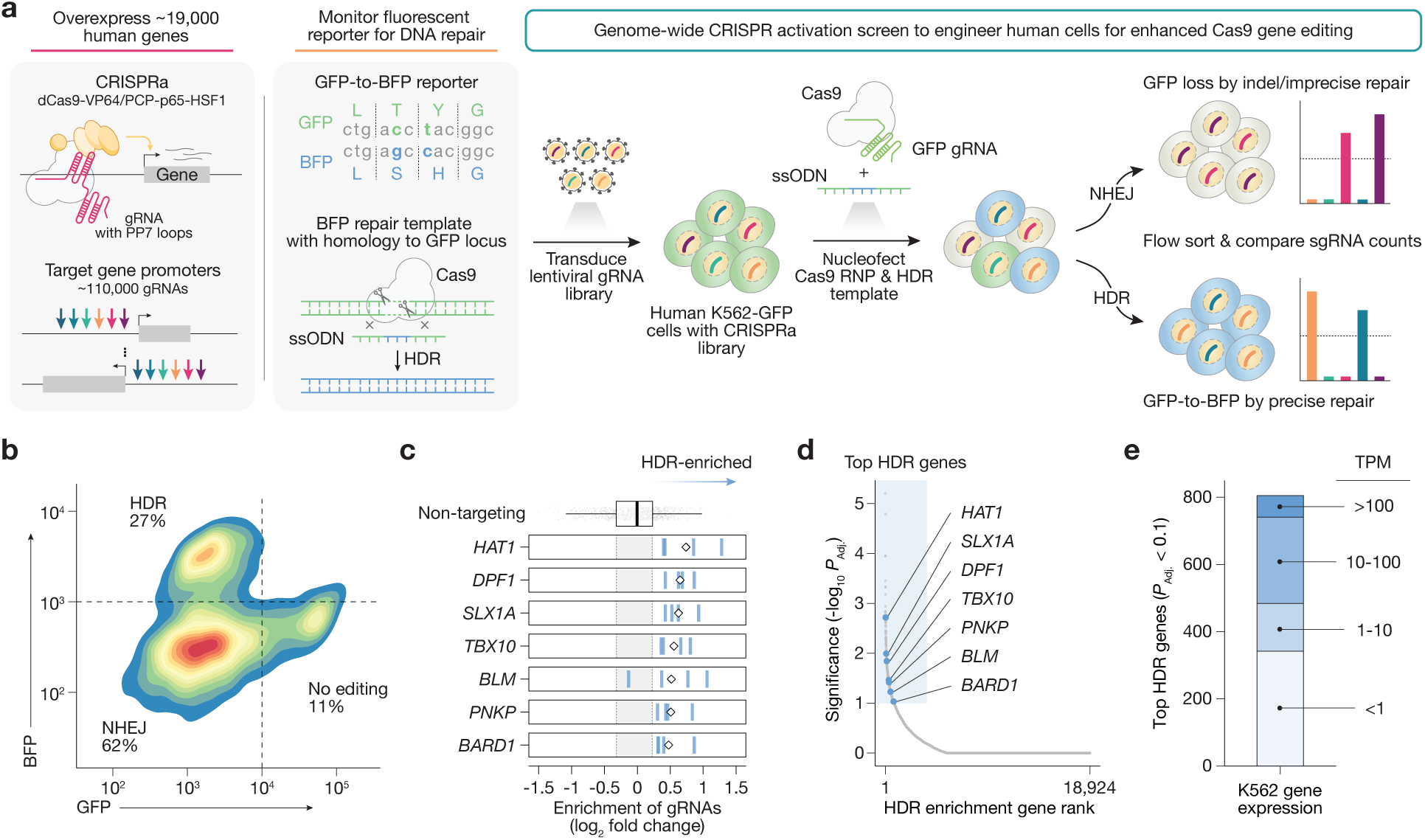
A genome-scale CRISPR activation (CRISPRa) screen identifies modulators of precise DNA repair in human cells. **a**, Human K562-EGFP cells were lentivirally transduced with CRISPRa (dCas9-VP64 + p65-HSF1) and a CRISPRa guide RNA (gRNA) library targeting ∼19,000 genes, selected, and nucleofected with a GFP-targeting Cas9 ribonucleoprotein complex and single-stranded repair template to convert GFP to BFP. Cells are flow sorted based on genome repair outcomes, and gRNA representation is computed in sorted bins for homology-directed repair (HDR) and non-homologous end-joining (NHEJ). **b**, Flow cytometry gating for repair outcomes (HDR or NHEJ) and quantification. **c**, Normalized enrichment in the HDR population for the four most enriched CRISPRa gRNAs targeting the indicated genes. The fold-change of each gRNAs is shown as blue lines and the average is indicated by the diamond. **d**, Gene ranks and significance for HDR enrichment with robust rank aggregation (RRA) using the four most enriched gRNAs. The highlighted region indicates genes with FDR-adjusted RRA *P*_adj_ < 0.1. **e**, RNA expression in K562 cells of the top enriched HDR genes (FDR *P*_adj_ < 0.1).

We first established a monoclonal human K562 cell line that stably co-expresses the CRISPRa effector dCas9-VP64^32^ and a single copy of a destabilized fluorescent *EGFP* reporter gene. We lentivirally transduced these cells with a genome-wide CRISPR library targeting the transcription start sites of ∼19,000 human genes with six guide RNAs (gRNAs) per gene^33^. This CRISPR library was optimized for on-target activity and co-expressed additional activators (PCP-p65-HSF1) that bind to the modified gRNA scaffold to maximize gene activation. We transduced cells with ∼113,000 CRISPRa gRNAs at a low multiplicity of infection — to ensure that most cells received only one gRNA — and then selected for transduced cells with puromycin.

After allowing 10 days for selection and target protein expression, we nucleofected the cells with a Cas9 ribonucleoprotein (RNP) targeting the GFP locus and a single-stranded DNA repair template to change GFP to BFP. After five days, we sorted BFP-positive (indicative of precise repair via HDR) and non-fluorescent (indicative of NHEJ repair) populations using fluorescence-activated cell sorting (**Fig. 1b**, **Supplementary Fig. 1a**).

Next, we extracted genomic DNA and performed amplicon sequencing to quantify CRISPRa gRNA abundance in the HDR and NHEJ groups (**Supplementary Table 1**). In each group, we aggregated the fold-change (compared to pre-sort cells) of different gRNAs targeting the same gene using robust rank aggregation (RRA) to identify genes with consistently enriched or depleted gRNAs (**Fig. 1c,d**, **Supplementary Table 2**).

In total, we found 805 genes that were enriched in the HDR population (*P_adj_* < 0.1) (**Fig. 1d**). We noticed that the top enriched genes in the HDR population included many known modulators of DNA repair (e.g. *SLX1A*^34^, *BARD1*^35^, *BLM*^36^, *PARP10*^37,38^, *SWSAP1*^39^, *RFC3*^40^) and also several other genes that have not previously been associated with DNA repair (e.g. *TBX10*, *ZNF146, ADH4, FOXO3*). Furthermore, we observed that more than half of highly enriched genes in the HDR population were expressed either lowly (1 to 10 transcripts per million [TPM]) or not at all (less than 1 TPM) in K562 cells (**Fig. 1e**). This highlights a key advantage of our gain-of-function approach: the ability to functionally characterize all human genes — including non-expressed genes — and their impact on protein function^41^. The same analysis was performed on the NHEJ screen and similarly identified known and novel modulators of DNA repair, although these genes were not of immediate interest for our protein design task (**Supplementary Fig. 1b-d**). In summary, our function-first, proteome-wide screen identified hundreds of proteins that modulate the balance between NHEJ and HDR. Thus, we aimed to use this functional catalog for the rational design of novel genome editors.

### Rational protein design for precise DNA repair

We reasoned that fusing top-ranked proteins from the functional HDR screen to Cas9 could further enhance HDR by localizing repair-promoting proteins at the locus to be edited (**Fig. 2a**). To nominate proteins for Cas9 fusion, we prioritized nuclear proteins with DNA-binding domains^42–44^ and selected 14 proteins ranking in the top 1% of our CRISPRa screen. We also included 6 proteins enriched in an independent overexpression screen for improved HDR^46^. This set included both genes with known roles in HDR (e.g. *BARD1*, *BLM*, and *RAD18*)^21,47^ and those without any previously described role (e.g. *ZNF146*, *TBX10*). In total, we cloned 20 full-length candidates as C-terminal fusions to Cas9 with an XTEN linker and named these fusion proteins Targeted Repair fUsion Editors (TruEditors) (**Supplementary Table 3**).

**Figure 2.**
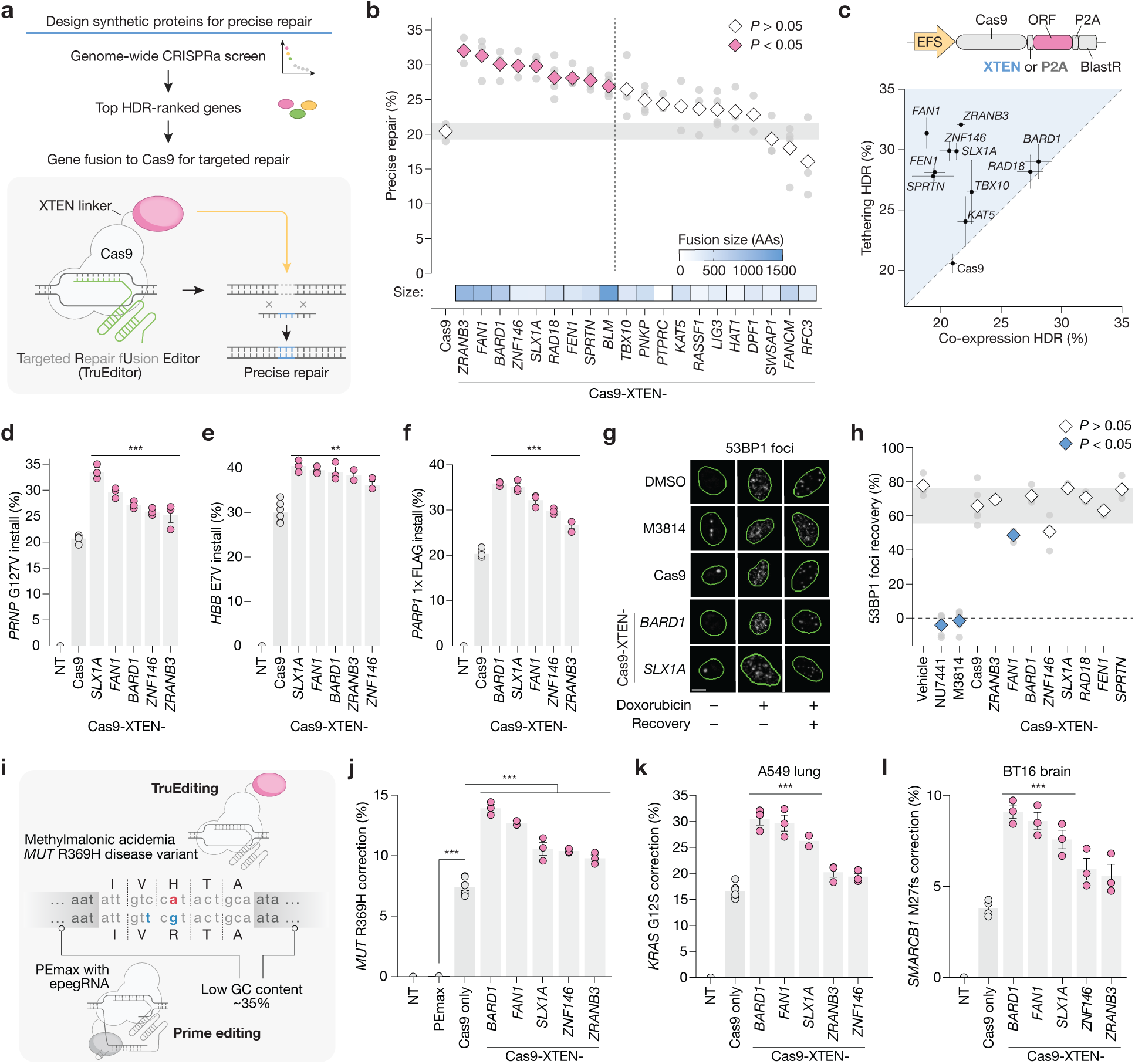
Rational design of TruEditors for precise gene editing. **a**, Candidates for protein fusion with Cas9 nuclease for improved precise gene editing. **b**, Precise repair (GFP-to-BFP conversion) for 20 TruEditors as measured by flow cytometry. TruEditor constructs were nucleofected into human kidney HEK293-EGFP cells with a GFP-targeting gRNA and a single-stranded repair template encoding BFP-specific edits. Points represent three independent transfection replicates and the diamond indicates the mean. Grey rectangle represents the 95% confidence interval for the Cas9 control. The heatmap shows the length of fused proteins. **c**, Comparison of TruEditors with significant increases in precise repair as direct Cas9 fusions (XTEN linker) versus co-expression (2A cleavage). Points represent the mean of three replicates with SEM error bars. **d**, Precise install of the protective G127V PRNP variant in HEK293 cells. **e**, Precise install of the sickle cell disease-causative HBB E7V variant in HEK293 cells. **f**, Precise knock-in of FLAG protein affinity tag at the N terminus of the *PARP1* gene in HEK293 cells. **g**, Representative immunofluorescent images of nuclear 53BP1 foci in human lung A549 cells expressing the indicated gene editor or treated with the NHEJ inhibitor M3814 (2 μM) or vehicle (DMSO) control. Cells were either treated for 4 hours with vehicle, doxorubicin (0.5 μM), or doxorubicin and then allowed to recover for 24 hours post-washout. Scale bar, 10 μm. **h**, Recovery of nuclear 53BP1 foci after doxorubicin treatment in A549 cells. **i**, Comparison of TruEditors and prime editing to correct the *MUT* R369H variant in HEK293 cells. **j**, Precise correction of *MUT* R369H variant in HEK293 cells. **k**, Precise correction of the lung cancer driver *KRAS* G12S mutation in A549 cells. **l**, Precise correction of the atypical teratoid/rhabdoid tumor (ATRT) driver *SMARCB1* M27fs mutation in human brain BT16 cells. For all comparisons, experiments had at least 3 independent replicates, and significance was determined using a one-way ANOVA. Post-hoc pairwise comparisons were conducted using Tukey’s HSD test (\**P* < 0.05, \*\**P* < 0.01, \*\*\**P* < 0.001).

We tested each TruEditor using the same GFP-to-BFP reporter system and, after plasmid and repair template nucleofection, we assessed editing outcomes by flow cytometry (**Fig. 2b, Supplementary Table 4**). We identified 9 TruEditors that significantly increased precise repair compared to Cas9 alone (one-way ANOVA *P* < 10^-16^, *F* = 77.1). Top-performing precise editors included fusions with key mediators of HDR, such as BRCA Associated Ring Domain 1 (BARD1). BARD1 recognizes H2A lysine 15 ubiquitination at DNA damage sites to recruit BRCA1 and direct HDR by antagonizing 53BP1^35,48^. We also validated several TruEditors with proteins that have not previously been implicated in HDR. For example, the ZNF146-based TruEditor yielded a ∼1.5-fold increase in HDR compared to Cas9 alone.

To test our hypothesis that localizing these proteins to the targeted locus is important for enhancing precise repair, we replaced the XTEN linker with a ribosomal-skipping P2A peptide that expresses Cas9 and the protein modulator separately while preserving equal stoichiometry^49,50^. We inserted the P2A sequence between Cas9 and 12 of the top-performing TruEditors that we tested previously (**Fig. 2c**). Only 2 of the 12 candidates (*BARD1* and *RAD18*) had similar improvements in HDR rates when the proteins were expressed separately or directly tethered to Cas9, highlighting the importance of localizing the protein modulators directly at the repair site (one-way ANOVA *P* < 10^-8^, *F* = 13.6).

Next, we aimed to assess whether editing improvements extend beyond our transgene reporter to four distinct endogenous genomic loci: installing a protective variant (G127V) in the prion disease protein *PRNP*^51^, installing the sickle-cell disease variant (E6V) in the beta-globin gene *HBB*^52^, and inserting in-frame two different affinity tags (23 and 69 amino acids in length) in the poly [ADP-ribose] polymerase 1 (*PARP1)* gene, an important drug target in breast and ovarian cancers (**Supplementary Table 4**). Using amplicon sequencing, we found in all cases that TruEditors improved precise editing over Cas9 alone, achieving precise editing rates between ∼25-40% (**Fig. 2d-f, Supplementary Fig. 2b**). To confirm tagging of *PARP1*, we used immunofluorescence (IF) to detect 1X- and 3X-FLAG tags inserted at the *PARP1* locus and found that the top TruEditor improved knock-in by 1.8-fold over Cas9 alone (**Supplementary Fig. 2c**).

To understand if TruEditor expression might alter DNA damages responses, we measured changes in double-strand breaks (γH2AX and 53BP1 foci) after inducing damage with doxorubicin (**Supplementary Fig. 2d,e**). For example, global DNA repair defects are a known problem with small molecule NHEJ inhibitors that improve HDR^30^: In human A549 cells, NHEJ inhibitors NU7441 and M3814 led to an increase in 53BP1 foci post-doxorubicin damage, whereas cells treated with vehicle had a more than 4-fold decrease in foci. In contrast to NHEJ inhibitors, we found that nearly all TruEditors assayed showed no impairment in DNA repair, highlighting a potential safety advantage over current methods to increase HDR (one-way ANOVA *P* < 10^-15^, *F* = 113) (**Fig. 2g,h, Supplementary Fig. 2f,g**).

We also tested if TruEditors can correct mutations that cause severe inborn genetic diseases that are challenging for other precise editing methods. Prime editing, for example, is inefficient at sites with low GC content^53^, such as the locus surrounding the methylmalonyl-CoA isomerase *MUT* R369H variant, which causes methylmalonic acidemia (MMA). This locus has a low GC content (∼35% in the surrounding 100 bp window) and other groups have reported that it cannot be corrected using prime editing^54^. We first attempted prime editing with an optimized system (PEmax with an epegRNA^55^) for the GFP transgene (64% GC content) and observed ∼29% BFP conversion (**Supplementary Fig. 2h**). Next, we attempted to fix the *MUT* R369H mutation using the same strategy, but this approach resulted in very low correction (< 0.1%), similar to previous observations^54^ (**Fig. 2i,j, Supplementary Fig. 2i,j**). In parallel, we delivered several TruEditors for precise editing and achieved correction rates of 10 - 15% at this locus. This was a significant improvement over Cas9 alone (up to ∼1.8-fold) for correcting the disease-causative allele (one-way ANOVA *P* < 10^-13^, *F* = 107.3).

To evaluate if TruEditor improvements apply to different cell types, we edited two additional disease-relevant loci. In A549 lung cancer cells, top-performing TruEditors, such as fusions with BARD1, FAN1 or SLX1A, nearly doubled the correction rate of a *KRAS* driver mutation (G12S) compared to Cas9 alone (**Fig. 2k**). These editors also more than doubled the correction rate of a *SMARCB1* frameshift deletion (M27fs) in BT16 brain tumor cells (**Fig. 2l**). Altogether, we designed and tested several synthetic gene editors by fusing top hits from the proteome-scale activation screen to the programmable Cas9 enzyme. These editors improved precise repair across diverse loci and cell types, outperformed Cas9 alone and corrected sites that cannot be edited using current state-of-the-art methods.

### Minimal TruEditors for precise gene editing

Since TruEditor fusion proteins span a range of sizes — from a few hundred residues to over a thousand residues — we wondered whether specific functional domains of these proteins might be sufficient for improved precise editing, yielding more compact TruEditors. We identified all annotated^56^ protein domains for the top 10 TruEditors and cloned 24 of these as minimal Cas9 fusions (**Fig. 3a,b, Supplementary Fig. 3a**). Of the 24 domain-only editors tested in the GFP-to-BFP reporter assay, three significantly improved HDR over Cas9 alone (one-way ANOVA *P* < 10^-15^, *F* = 33) (**Fig. 3c, Supplementary Fig. 3b**). For instance, the BARD1 Zinc-finger, RING-type domain (BARD1_32-117_) improved editing ∼1.4-fold and the KAT5 histone acetyltransferase domain (KAT5_174-451_) showed a ∼1.3-fold improvement over Cas9.

**Figure 3.**
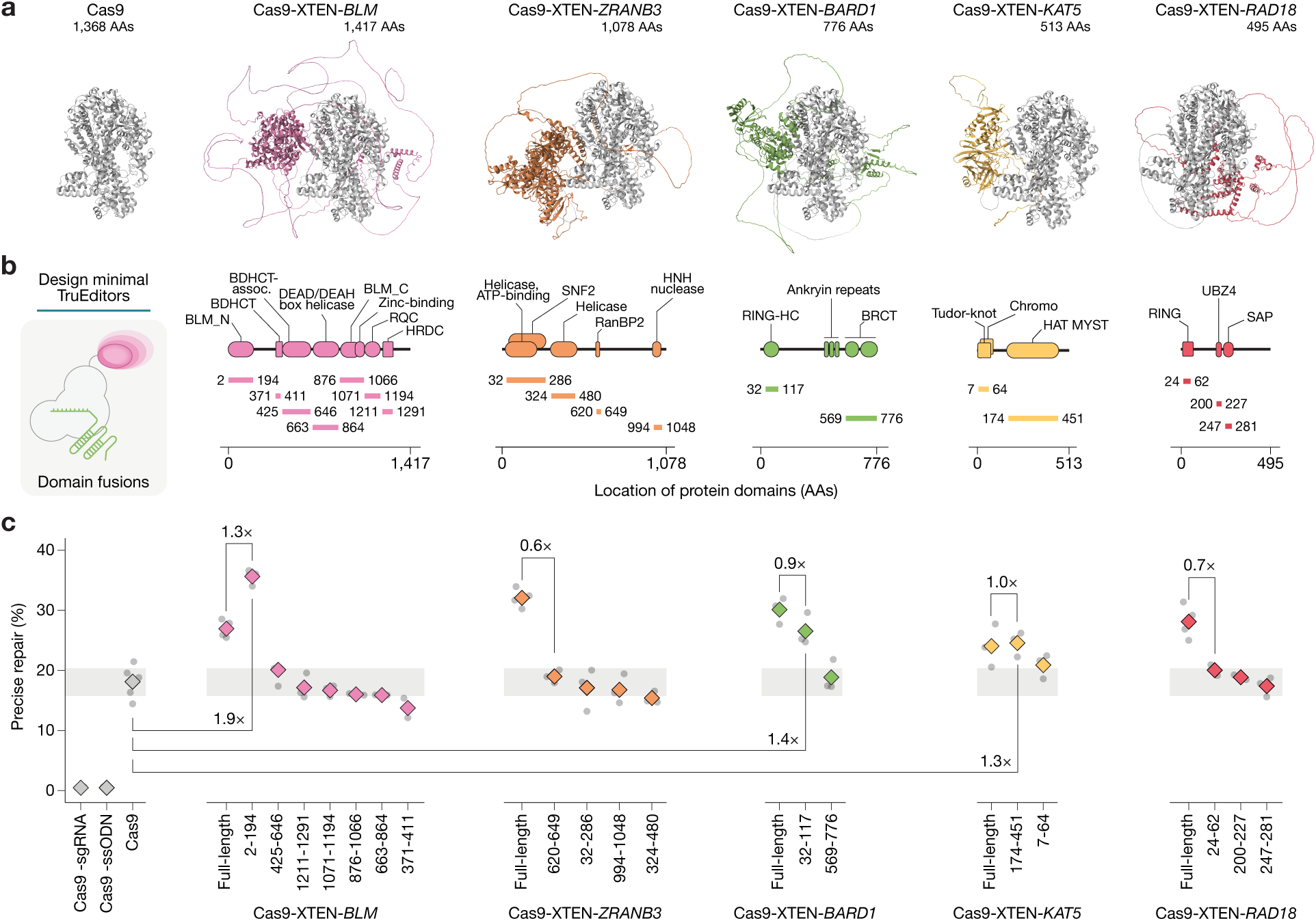
Development of minimal TruEditors by replacing full-length proteins with small protein domains. **a**, AlphaFold3 structure predictions for the indicated, full-length Cas9 fusion TruEditors. **b**, Linear representation of annotated protein domains from the indicated proteins. Cloned domains or combinations of domains are indicated as horizontal lines and their lengths given in amino acids. **c**, Precise repair (GFP-to-BFP editing) rates for the indicated minimal or full-length TruEditors. TruEditor constructs were nucleofected into HEK293-EGFP cells with a GFP-targeting gRNA and a single-stranded repair template encoding BFP-specific edits. Points represent three independent transfection replicates, and the diamond indicates the mean. Grey rectangle represents the 95% confidence interval for the Cas9 control.

The top-performing fusion editor was a minimal domain from the N-terminus of the Bloom Syndrome RecQ-like Helicase BLM (BLM*_2_*_-194_). This minimal editor improved HDR rates ∼1.9-fold over Cas9 and outperformed the full-length Cas9-BLM TruEditor by ∼1.3-fold (**Fig. 3c**). We also confirmed that its activity required direct tethering to Cas9 (**Supplementary Fig. 3c**). We then validated its broad applicability across seven different precise edits in three distinct cell models, including the five edits previously tested, and insertion of an affinity tag to the *ZNHIT1* gene and correction of the KRAS G13D mutation in MDA-MB-231 breast cancer cells (**Supplementary Fig. 3d-e**). In every context, the minimal Cas9-BLM_2-194_ TruEditor significantly improved precise editing rates over Cas9 alone.

Notably, the top-performing domain fusions — BLM_2-194_ and BARD1_32-117_ — are both known to mediate protein-protein interactions within HDR complexes. Specifically, the BLM domain is essential for forming the BLM:TOP3A:RMI1/2 (BTR) complex^57,58^, while the BARD1 domain is required for the BARD1:BRCA1 heterodimer formation^59,60^. Thus, we hypothesized that TruEditors may function by recruiting distinct endogenous proteins to the edit site.

### TruEditors coordinate distinct endogenous DNA repair proteins

To characterize the protein-protein interaction (PPI) networks of the top six performing TruEditors, we performed affinity purification-mass spectrometry (AP-MS) (**Fig. 4a**). We expressed the TruEditors as a FLAG-tagged bait protein in HEK293FT cells and then purified their interacting protein complexes via FLAG immunoprecipitation. These complexes were then analyzed by mass spectrometry. Replicates for each TruEditor bait showed very high peptide spectral match (PSM) correlation (*r* > 0.88, **Fig. 4b**).

**Figure 4.**
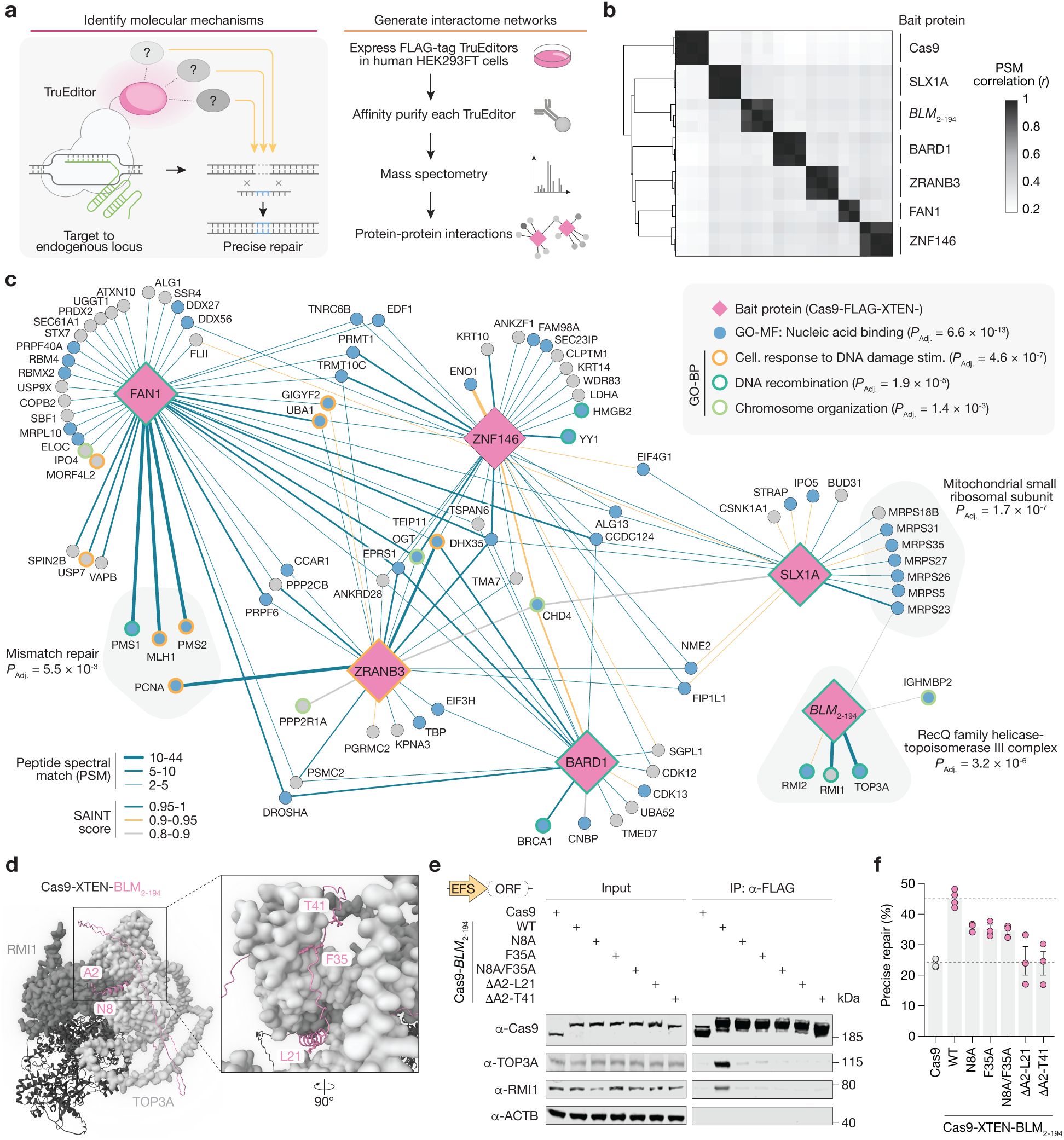
TruEditors coordinate shared and distinct groups of proteins with roles in DNA repair. **a**, Affinity proteomics (AP) to identify protein-protein interactomes for different TruEditors. FLAG-tagged TruEditors were immunoprecipitated from HEK293 cells after 24 hours and interacting proteins were identified on a Orbitrap Eclipse Tribrid mass spectrometer. **b**, Pearson correlation of peptide spectral match (PSM) counts (*n* = 2 - 3 biological replicates per TruEditor). **c**, Protein-protein interaction networks for TruEditors and their significant interactors (average PSM fold-change over Cas9 alone > 10 and *P*_adj_ < 0.05). Edge thickness (average PSM) and color (significance analysis of interactome [SAINT] score^103^) correspond to the strength of the connection. Nodes are colored by Gene Ontology category. **d**, Ternary complex of the Cas9-*BLM*_2-194_ TruEditor with AP-validated interactors *TOP3A* and *RMI1* as predicted by AlphaFold3. Key *BLM*_2-194_ interaction residues at the interface with *TOP3A* (*light grey*) and *RMI1* (*medium grey*) are highlighted in *pink*. **e**, Co-immunoprecipitation of FLAG-tagged Cas9 only, Cas9-*BLM*_2-194_ and different Cas9-*BLM*_2-194_ mutants (N8A, F35A, N8A/F35A, ΔA2-L21, ΔA2-T41) in HEK293 cells. **f**, Precise editing rates (GFP-to-BFP editing) for Cas9 only, Cas9-*BLM*_2-194_ and Cas9-*BLM*_2-194_ mutants in HEK293-GFP cells as measured by flow cytometry (*n* = 3 biological replicates).

In total, we identified 146 high-confidence PPIs involving 91 individual proteins (*p_adj_* < 0.05, fold-change over Cas9 alone > 10 and PSM > 2.5). These interacting proteins were significantly enriched in Gene Ontology (GO) terms related to DNA repair, such as cellular response to DNA damage stimulus, DNA recombination, and chromosome organization (**Fig. 4c, Supplementary Fig. 4a, Supplementary Table 5**). In addition, we found that over half of the interacting proteins (52 out of 89) were enriched for nucleic acid binding (*P_adj_* = 6.6 × 10^-13^). Notably, we discovered both known and novel PPIs. For instance, we captured well-known complexes like the BARD1:BRCA1 complex^48^, the BTR complex (BLM:TOP3A:RMI1/RMI2)^57^, and the FAN1:MLH1:PMS1:PMS2 complex^61^ (**Fig. 4c, Supplementary Fig. 4d-f, Supplementary Movies 1-3**). Beyond these well-studied complexes, we also discovered novel PPIs. Most notably, we identified an interaction between ZNF146 and the known DNA damage response factor YY1 (**Fig. 4c, Supplementary Fig. 4g, Supplementary Movie 4**). YY1 is an early DNA damage response factor, which stimulates PARP1 enzymatic activity through direct binding at sites of DNA damage^62,63^. Given the limited literature suggesting any direct link between ZNF146 and DNA repair^64–66^, the PPI with YY1 offers a potential mechanism for the improved editing rates of the ZNF146-based TruEditor.

Interestingly, we also identified 7 common PPIs across most TruEditors, including CHD4, DDX56, DHX35 and RBM5, which were enriched in 5 of the 6 baits (**Fig. 4c, Supplementary Fig. 4b,c**). These common proteins suggest a unified mechanism connecting distinct TruEditors. For instance, CHD4, a helicase and component of the NuRD chromatin remodeling complex, plays an important role in relaxing DNA around DSB sites and regulating HDR through coordination with EP300 and BRIT1^67–69^.

To investigate how the top-performing TruEditor (Cas9-BLM_2-194_) improves HDR, we used AlphaFold3^70^ to model the PPIs identified by AP-MS. We predicted the interacting residues where the BLM_2-194_ domain binds TOP3A and RMI1 (**Fig. 4d, Supplementary Movie 5**). We identified two structured regions of high prediction confidence (pLDDT > 50) corresponding to an alpha helix and beta sheet near the N-terminus (residues 7 to 38). The alpha helix N7-N22 facilitates interaction between BLM_2-194_ and TOP3A, while the beta sheet G33-K38 interacts with RMI1, based on residue proximity and confidence scores from AlphaBridge^71^ (**Supplementary Fig. 4h**).

To test these predicted interactions, we created alanine point mutants at key residues (N8A, F35A) and truncation mutants lacking the predicted binding interfaces (residues 1-20 or 1-40) (**Supplementary Fig. 4i**). Co-immunoprecipitation (Co-IP) experiments confirmed that these mutations disrupted binding to TOP3A and RMI1 (**Fig. 4e**). Surprisingly, individual point mutations were sufficient to disrupt the entire complex, suggesting all three proteins are required for stable interaction. We also tested whether these mutants with disrupted binding interactions were able to enhance precise repair using the HDR reporter assay. All point and truncation mutants showed significantly lower HDR rates than the wild-type BLM_2-194_ TruEditor (one-way ANOVA *P* < 10^-5^, *F* = 13.5, **Fig. 4f**). These results demonstrate that the BLM_2-194_ domain recruits endogenous proteins that are required for improving HDR.

### Precise editing in human embryonic stem and primary T cells via mRNA delivery

Given the improvements in precise repair and the versatility of TruEditors for site-specific corrections, we next sought to optimize their delivery for therapeutic applications. Recently, mRNA has been widely used as a vehicle for gene editing proteins due to its low immunogenicity^72–75^. Thus, we synthesized fully modified (5′ Cap-1 structure and internal pseudouridine (Ψ)-substitutions) TruEditor mRNAs with engineered untranslated regions (UTRs)^76^ for robust expression in human embryonic stem cells (hESC) and primary T cells (**Fig. 5a**).

**Figure 5.**
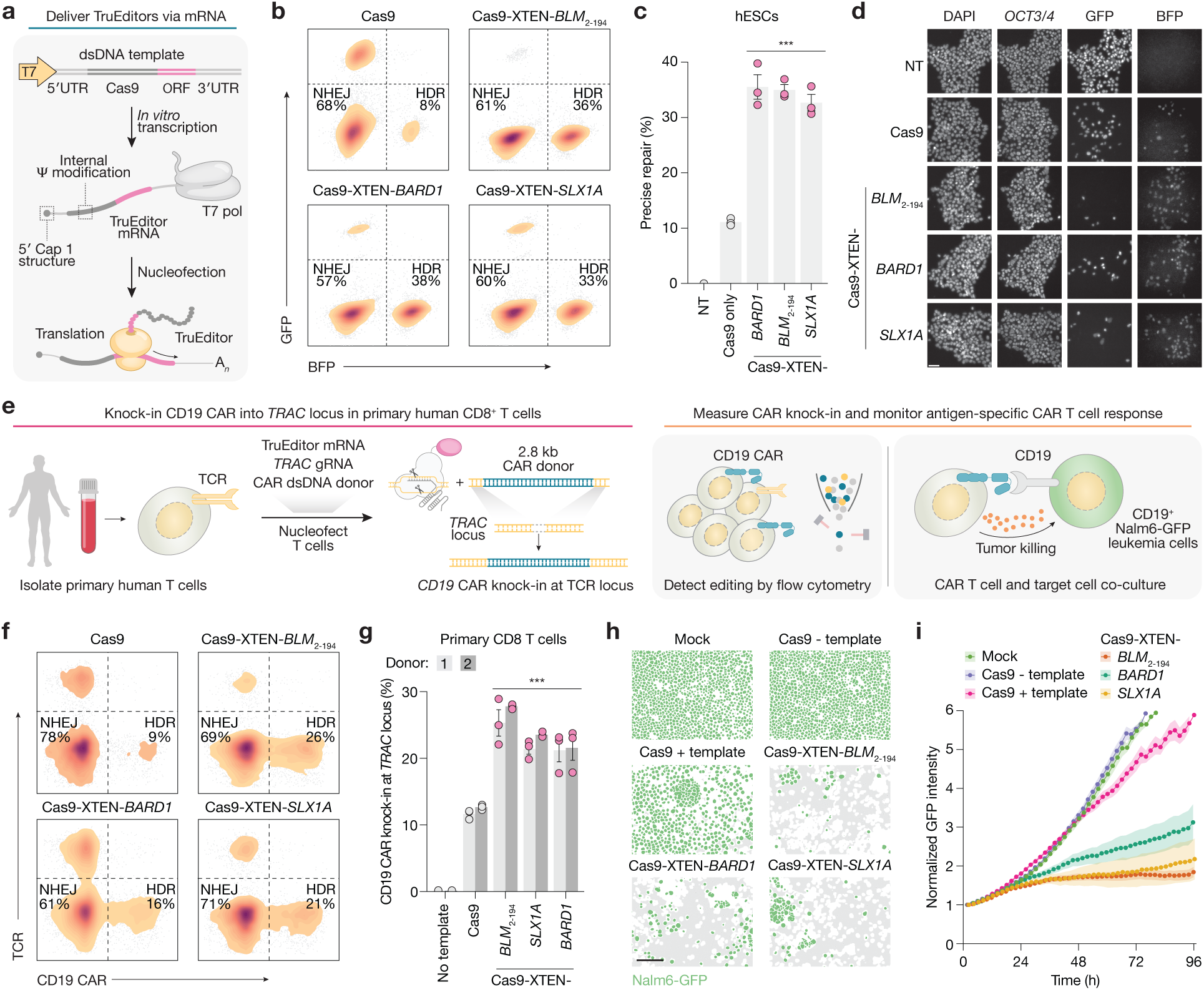
Messenger RNA (mRNA) delivery of TruEditors in stem cells and primary human T cells improves precise gene editing. **a**, In vitro transcription to generate TruEditor mRNAs for high efficiency and low immunogenicity delivery. **b**, Representative flow cytometry plots from HUES66-GFP human embryonic stem cells to measure precise gene editing via GFP-to-BFP conversion. **c**, Precise gene editing in HUES66-GFP human embryonic stem cells nucleofected with TruEditor mRNA, GFP-targeting gRNA, and a single-stranded repair template encoding BFP-specific edits (*n* = 3 biological replicates). **d**, Representative native fluorescent protein and immunofluorescence images from HUES66 human embryonic stem cells. Scale bar, 50 μm. **e**, Knock-in of a 2.8 kb anti-CD19 chimeric antigen receptor (CAR) into the *TRAC* locus in primary CD8+ human T cells and quantification of editing and tumor killing (*n* = 2 human donors). **f**, Representative flow cytometry plots to measure CAR knock in and TCR loss. **g**, Quantification of CAR knock-in and TCR loss in primary human T cells by flow cytometry (*n* = 2 human donors with 3 nucleofection replicates each) **h**, Representative images from primary CD8+ human T cells edited as in **f,g** after 96 hours in co-culture with CD19+ Nalm6-GFP leukemia cells (effector:target [E:T] ratio of 1:2). Scale bar, 100 μm. **i**, Normalized GFP intensity for CD19+ Nalm6-GFP leukemia cells in co-culture with primary CD8+ human T cells edited as in **f,g** (*n* = 24 images per condition with 4 images per nucleofection and 3 nucleofections for each of 2 human donors). For all comparisons, statistical significance was determined using a one-way ANOVA with Tukey’s HSD post hoc test (\**P* < 0.05, \*\**P* < 0.01, \*\*\**P* < 0.001).

We first tested TruEditor mRNAs in HUES66 hESCs engineered with a GFP-to-BFP reporter. We observed ∼3-fold improvements in precise editing with BARD1, BLM_2-194_, and SLX1A TruEditors compared to Cas9 (**Fig. 5b,c, Supplementary Fig. 5a-c**). This was the largest editing improvement observed at any locus or cell line tested, highlighting the potential of TruEditors delivered as mRNA to improve precise editing in stem cells, which historically have been more difficult to edit than other cell lines due to their intact p53 function^77^. We also confirmed that editing did not alter pluripotency, as cells retained strong expression of *OCT3/4,* comparable to untreated controls (**Fig. 5d**).

A key advantage of HDR editing over both prime and base editing is the ability to install kilobase-scale, in-frame knock-ins^78,79^. This strategy has been used for engineering chimeric antigen receptor (CAR) T cells through the knock-in of CAR genes into the *TRAC* locus, which encodes the constant region of the T cell receptor (TCR) alpha chain^80^. This knock-in approach has been shown to be superior to viral transduction or transposon insertion, as it leverages endogenous promoters and enhancers to delay T cell exhaustion^81,82^. Knock out of the endogenous TCR also prevents donor T cells from mounting an alloreactive response against the recipient (graft-versus-host disease), enabling allogeneic cell therapies using donor T cells to treat many cancer patients without HLA matching. We therefore sought to understand whether TruEditors could improve knock-in efficiency of a CAR in primary human T cells.

First, we generated a 2.8 kb double-stranded HDR template for a CD19 CAR, flanked with Cas9 target sequences to enhance nuclear import^80^. We delivered this template, along with the TruEditor mRNA and a gRNA targeting the *TRAC* locus, into primary human CD8⁺ T cells from two healthy donors. After five days, we used flow cytometry to measure TCR knockout and CAR knock-in rates (**Fig. 5e, Supplementary Fig. 5d,e**). TruEditors improved the knock-in efficiency by more than 2-fold over Cas9 alone (**Fig. 5f,g, Supplementary Fig. 5f**).

Next, we assessed the therapeutic function of these engineered CAR-T cells. We co-cultured CAR-T cells with CD19-positive leukemia cells (Nalm6-GFP) and monitored cancer cell killing via live-cell imaging. The TruEditor-engineered CAR-T cells were significantly more effective at killing the leukemia cells than those engineered with Cas9 alone (one-way ANOVA *P* < 10^-16^, *F* = 72.77) (**Fig. 5h,i, Supplementary Fig. 5g,h**). Together, these results demonstrate the high efficiency and safety of TruEditor mRNA delivery in human stem cells and primary T cells and establishes TruEditors as a versatile and potent modality for engineering next-generation cell therapies.

## Discussion

Our results demonstrate that overexpression of ∼18,000 human proteins can inform and guide the rational design of proteins with specific functions. Here, we capitalized on a genome-scale overexpression screen to identify positive modulators of precise gene editing and design Cas9 fusion proteins. These fusion proteins (TruEditors) robustly improve precise editing rates across a diverse set of cells, genomic loci and types of edits. We demonstrate that TruEditors benefit from direct Cas9 tethering to bring them to the edit site, that they recruit specific proteins that regulate DNA repair, and that they are efficient at small (1-3 bp), medium (20-100 bp), and large (>2 kb) edits, including in primary human cells and at sites where other editing modalities, such as prime editing, do not work. We anticipate TruEditors will also be useful in combination with other chemogenetic perturbations that can improve HDR^22,23,80^ and that similar genome-scale protein engineering approaches could be applied to other editing modalities such as prime or base editing.

Given the multiple clinical trials underway using somatic cell editing to correct inborn genetic disorders, a key concern is the potential immunogenicity of any novel Cas9 fusion. Compared to many other CRISPR editors that rely on DNA-modifying enzymes from bacterial and non-human sources, an inherent advantage of using human proteins/domains, as we did here, is reduced potential immunogenicity of TruEditors. However, this needs to be tested rigorously before clinical application. Another important consideration is that, given the diversity of mechanisms identified for different TruEditors, these proteins may perform variably depending on the cell or tissue type. For instance, we observe that for most cell types tested in this study, Cas9-BARD1 is the top-performing TruEditor; however, for primary human T-cells, Cas9-BLM_2-194_ shows the highest rates of precise editing. This may be due to variable expression of specific co-factors that are required. Importantly, the diversity of TruEditors also creates an opportunity for further context-specific optimization, which is more challenging for precise editing approaches that rely on a single Cas9 enzyme fusion.

A surprising finding is that, in certain cases, subregions or domains of proteins can drive similar improvements in precise genome editing as the full-length human protein. In this work we identify the *N*-terminal domain of BLM as a potent and compact modulator of HDR when tethered to Cas9. We expect that additional short domains and protein binders exist which can similarly modulate DNA repair, and that discovery of such compact sequences will enable combinatorial protein fusions with even greater potential. Since the space of possible combinations for protein or domain fusions is enormous, the genome-scale overexpression approach can also guide choices to reduce the combinatorial protein design space, resulting in faster and more cost-effective campaigns than *de novo* protein design approaches with limited measurements of protein function.

Overall, this study provides the first genome-scale functional map of positive modulators of precise HDR, and applies insights gained to develop TruEditors, a novel class of precise gene editors. We envision that similar function-guided genome-scale overexpression can — perhaps in combination with emerging deep learning frameworks — enable the design of new proteins to improve any selectable phenotype.

## Methods

### Cell line culture and maintenance

HEK293FT cells were acquired from Thermo Fisher Scientific (Thermo R70007) and cultured in DMEM (Cytiva SH30022) supplemented with 10% Serum Plus II (Sigma-Aldrich 14009C) or HyClone FetalClone III serum (Cytiva SH30109) by volume (D10 media). BT16 cells were gifts from P. Houghton (University of Texas Health, San Antonio), R. Hashizume (University of Alabama, Birmingham) and C. Roberts (St. Jude Children’s Research Hospital). K562, MDA-MB-231, and A549 cell lines were obtained from American Type Culture Collection (ATCC). BT16, A549, and MDA-MB-231 cells were also cultured in D10 media. K562 cells were cultured in I10 media: IMDM (Thermo 31980097) plus 10% Serum Plus II or FetalClone III serum.

HUES66 human embryonic stem cells (hESCs) were acquired from the Harvard Stem Cell Institute and cultured on LDEV-free Geltrex-coated plates (Thermo A1413201) in Essential 8 (E8) medium (Thermo A1517001). E8 media was supplemented with CEPT cocktail (Tocris 7991) for 18-24 hours after hESC passaging and then removed.

All cell lines were maintained at 37 °C and 5% CO_2_ with regular passaging and regular mycoplasma testing using the MycoAlert Plus Mycoplasma Detection Kit (Lonza LT07-701).

### Lentiviral transduction and transposon integration for cell line transgenesis

Lentivirus was produced by transfecting HEK293FT producer cells plated in 6-well dishes with 1 μg of a transfer plasmid encoding the desired expression vector alongside 0.8 μg of the packaging plasmid psPAX2 (Addgene 12260) and 0.55 μg of the envelope plasmid pMD2.G (Addgene 12259), using 5.5 μL linear polyethylenimine (PEI, MW 25,000; Polysciences 23966). Transfer plasmids included EGFP-d2PEST-2A-Hygro plasmid (Addgene 138152) to generate GFP expression lines, dCas9_VP64-Blast (Addgene 61425) for K562-GFP-CRISPRa, lentiCas9-Blast (Addgene 52962) for A549-Cas9 and HEK293FT-Cas9 lines, and others cloned for this study (see *Plasmid design and cloning*). After 8 hours, transfection media was replaced with D10 supplemented with 1% bovine serum albumin (VWR AAJ65097-18). Viral supernatants were collected at 72 hours post-transfection, filtered through a 0.45 μm filter (Millipore SE1M003M00), aliquoted, and immediately stored at −80 °C for later use.

Single-copy monoclonal EGFP cells were used for GFP-to-BFP editing experiments: HEK293FT or K562 cells were transduced at low multiplicity of infection (MOI) with the EGFP-Hygro vector and selected with 50 μg/mL Hygromycin B (Thermo 10687010) for 10 days. Monoclonal GFP-expressing lines were established through dilution and clonal expansion in 96-well plates. To generate the K562-GFP-dCas9_VP64 line, GFP-positive K562 cells were further transduced with the dCas9_VP64-Blast lentiviral vector and selected with 10 μg/mL Blasticidin S (Thermo A1113902).

Polyclonal TruEditor- or Cas9-expressing A549 and HEK293 cell lines used in DNA damage and repair immunofluorescence assays were similarly generated by lentiviral transduction with editor-encoding vectors, followed by selection with 1 μg/mL puromycin (InvivoGen ant-pr-1) for at least 3 days. HEK293 cells expressing a non-targeting guide RNA (gRNA) used in AP experiments and HEK293 MUT exon 6 mutant cell lines used for R369H correction experiments were generated similarly and selected with 1 μg/mL puromycin for at least 3 days.

HUES66 hESCs lines were generated through piggyBac transposition. We used 250 ng total DNA in a 2:1 ratio of pPB-TO-oNgn2-puro-2A-GFP to SuperPiggyBac transposase. The day before transfection, 2 × 10⁵ cells were plated in 12-well Geltrex-coated plates in 1 mL Essential 8 medium (E8) + CEPT. The next morning, medium was replaced with 0.5 mL E8. Plasmids were diluted in 25 µL Opti-MEM I Reduced Serum Medium (Thermo 31985070) and mixed with 25 µL Opti-MEM containing 2 µL Lipofectamine Stem Transfection Reagent (Thermo STEM00001). After 10 minutes at room temperature, 50 µL transfection mix was added dropwise to each well. After 6 hours, 1 mL E8 was added. At 48 hours post-transfection, medium was replaced with E8 containing 1 µg/mL puromycin. Puromycin selection (1 µg/mL) was maintained for 5 days, after which cells were returned to standard culture conditions.

### Genome-scale CRISPR activation pooled screen: Lentiviral transduction and Cas9 RNP nucleofection

The Calabrese human CRISPR activation pooled libraries A and B (Addgene 1000000111) were used for the genome-scale overexpression screen. Briefly, 225 cm^2^ flasks of 80% confluent HEK293FT cells were transfected with 25 µg Calabrese library plasmid, 14 µg pMD2.G and 20 µg psPAX2 mixed in 2.5 mL OptiMEM and 175 µL polyethylenimine linear MW 25000 (1 mg/mL). At 6 hours after transfection, the media was exchanged for D10 supplemented with 1% bovine serum albumin. After 60 hours from transfection, viral supernatants were harvested and centrifuged at 3,000 rpm and 4 °C for 20 minutes and filtered using a 0.45 µm filter. The supernatant was then ultracentrifuged at 100,000 × g for 2 hours (Sorvall Lynx 6000) and the resulting pellet was resuspended overnight at 4 °C in PBS with 1% BSA.

Following lentiviral titration, 3 × 10^8^ K562-GFP-dCas9VP64 cells were transduced at an MOI of 0.3 and selected for 7 days with 2 µg/mL puromycin in order to allow overexpression to take effect. We nucleofected 75 million cells from the screen pool with a precomplexed GFP-targeting gRNA (IDT Alt-R CRISPR-Cas9 single-guide RNA [sgRNA]) and NLS-Cas9-NLS purified protein (Aldevron 9212) along with a BFP single-strand oligodeoxynucleotide (ssODN) (IDT Ultramer) using a Lonza 4D Nucleofector. To complex the gRNA with the Cas9, 60 µg of Cas9 protein was mixed with 10 µL 1× Cas9 buffer (GenScript), and, separately, 10 µL GFP gRNA (resuspended at 100uM) was mixed with 10 µL 1× Cas9 buffer. Then, the Cas9 and gRNA tubes were mixed, incubated at room temperature for 15 minutes, then added to 10 million human cells resuspended in 100 µL SF buffer (Lonza V4XC-2024), and nucleofected according to the manufacturer protocol. Following 5 days in culture, 200 million cells were sorted on Sony SH800S cell sorter into 3 populations: BFP+GFP-, GFP-BFP-, and GFP+BFP-, which were promptly lysed for gDNA extraction.

### Genome-scale CRISPR activation pooled screen: Genomic DNA extraction and library amplification

Screened cells were lysed using 12 mL of NK Lysis Buffer (50 mM Tris, 50 mM EDTA, 1% SDS, pH 8 per up to 100 million cells. 60 uL of 20 mg/mL Proteinase K (Qiagen 19133) was then added and each sample was incubated at 55 °C overnight. The next day, 60 uL of 20 mg/mL RNase A (Qiagen 19101) was added, and the samples were incubated at 37 °C for 30 minutes. Next, 4 mL of chilled 7.5 M ammonium acetate was added, and samples were vortexed then spun at 4,000 × g for 10 minutes. The supernatant was harvested and mixed with 12 mL isopropanol, then spun at 4,000xg for 10 minutes. Finally, DNA pellets were washed with 12 mL of 70% ethanol, spun, dried and pellets were resuspended with 0.2× TE buffer (Sigma 93302).

For the first PCR reaction of the library preparation, all gDNA available was used for each sample. 15 cycles of PCR 1 were performed using Taq-B polymerase (Enzymatics P7250L) in multiple reactions where each reaction contained up to 10 ug gDNA per 100 uL. PCR1 products for each sample were then pooled and used as input for PCR2 with barcoded primers. 12 PCR2 reactions for each sample were performed using 5 µL of the pooled PCR1 product per 100uL PCR2 reaction with Q5 polymerase (NEB M0491L). PCR2 products were pooled and normalized within each biological sample before combining uniquely barcoded separate biological samples. The pooled product was gel-purified from a 2% E-Gel (Thermo G402002) using the QiaQuick Gel Extraction kit (Qiagen 28704). The purified, pooled library was quantified with Tapestation 4200 (Agilent Technologies). Sequencing was performed on the NextSeq 550 instrument using the HighOutput Mode v2 with 75 bp paired-end reads (Illumina).

### Genome-scale CRISPR activation pooled screen: Read alignment and analysis

Sequencing reads were demultiplexed upon sequencing based on Illumina i7 barcodes present in PCR2 reverse primers using Illumina BaseSpace. Adaptor trimming was performed by treating the hU6 promoter sequence as an adaptor, using cutadapt v1.13 [`-e 0.2 -O 5 -m 20 -g TCTTGTGGAAAGGACGAAACACCG`]. Processed gRNA sequences were aligned to the Calabrese reference^83^ allowing for up to 1 mismatch using bowtie v1.1.2 [`-a–best–strata -v 1 -norc`]. Guide RNA counts were normalized to the total number of reads in each sample (**Supplementary Table 1**). Prior to computing gene ranks, gRNAs without sufficient representation in the presort (preselection) population were removed from analysis. Gene enrichments were computed using robust rank aggregation with the 4 most enriched gRNAs (minimum of 10 reads per gRNA). To determine genes in the top 1% of the CRISPRa screen, we used MAGeCK-RRA with all 6 gRNAs (minimum of 75 reads per gRNA).

K562 transcript gene expression data is from DepMap^85^ (release 24Q4). Protein localization data for all genes in the CRISPRa screen was taken from the Human Protein Atlas database (https://www.proteinatlas.org/)^42^, which uses experimental evidence (i.e. staining and microscopy) to determine intracellular localization; protein length in residues was taken from the UniProt database (https://www.uniprot.org/)^43^; and the presence of a DNA binding domain was predicted using DRNApred, which has previously been run on every native human protein (http://biomine.cs.vcu.edu/servers/DRNApred/)^44^.

### Plasmid design and cloning

Lentiviral Cas9 fusion vectors were cloned using T5 Exonuclease Dependent Assembly (TEDA)^86^. First, a parent vector was generated containing a XTEN linker with Esp3I cloning sites (IDT gBlock) TEDA cloned into a BamHI-digested lentiCas9-Blast backbone (Addgene 52962). Fusion ORFs were PCR-amplified from either complementary DNA (cDNA) or a library of human ORFs^87^ and TEDA cloned into the Esp3I-digested parent vector. The same strategy was used for the generation of lentiviral domain-only TruEditor constructs by amplifying specific domains from cloned, full-length TruEditor constructs.

To create cDNA libraries, RNA was extracted from 2 × 10^5^ HEK293FT cells lysed in 400 µL TRIZOL reagent (Thermo 15596018) using the Direct-zol RNA Microprep kit (Zymo R2062). Next, first strand synthesis was performed on 500 ng of the purified RNA product using the RevertAid First Strand cDNA Synthesis kit (Thermo EP0442). Finally, 20 µL of the cDNA product was diluted 1:5 in H20 and 2 µL of the resultant library were used to PCR genes for cloning with Q5 polymerase (NEB M0491L).

Co-expression TruEditor vectors for comparison against direct tethering (Fig. 2c) were generated similarly to direct fusion TruEditors: a parent vector containing a codon optimized, unique P2A sequence was synthesized (IDT gBlock) and cloned into a BamHI-digested lentiCas9-Blast backbone (Addgene 52962). Next, select fusion ORFs were amplified from previously cloned vectors and TEDA cloned into the parent vector.

All gRNA vectors were cloned by restriction cloning with annealed oligo pairs. Guide RNA top and bottom oligo pairs with designed overhangs were individually synthesized (IDT) and annealed, then assembled into an Esp3I-digested lentiGuideFE-Puro backbone (Addgene 170069) or into a BsaI- and HindIII- digested small, transfection vector (Addgene 174039) with addition annealed scaffold oligos using T4 ligation (NEB M0202L). pegRNA vectors were cloned according to Doman et al.^88^ using pU6-tmpknot-GG-acceptor (Addgene 174039).

TruEditor vectors for subsequent nucleofections of additional cell types (Fig. 2d-f,i-l) were cloned into smaller, transfection vectors using HiFi Gibson Assembly (NEB E2621L). First, an analogous parent vector was generated containing hspCas9, the XTEN linker, and a Esp3I cloning site by HiFi assembly of a PCR product from the parental vector into a NotI- and AgeI-digested CMV vector (Addgene 174820).

Cas9-BLM_2-194_ mutant vectors for AP (Fig. 4f) were generated by synthesizing the intended mutants (IDT gBlock) and HiFi Gibson Assembly. All TruEditor plasmids were endo-free midi-prepped (Qiagen 12945) and sequence verified by whole-plasmid sequencing.

Vectors for mRNA generation through in vitro transcription were generated by subcloning TruEditors into a pT7 vector (Addgene 214813) through PCR and HiFi Gibson Assembly. Inserts of entire TruEditor ORFs were amplified and the backbone vector was linearized through PCR prior to assembly. The pPB-TO-oNgn2-puro-2A-GFP plasmid was created by cloning GFP into iC13-PB-TO-oNgn2-BFP-puro (Addgene 204733, a gift from iPSC Neurodegenerative Disease Initiative [iNDI] & Michael Ward).

A *CD19* CAR-T donor vector was generated by subcloning an anti-*CD19* CAR (Addgene 181975) into a *TRAC* homology arm vector (Addgene 186127) using PCR to both linearize the backbone and generate the insert for HiFi Gibson assembly with 30 nt overhangs.

### Guide RNA and repair template design and delivery via nucleofection

The gRNAs for HDR editing experiments were designed using Benchling. We prioritized target sequences which were proximal to the edit site (within 1-20 bps—ideally within 1-10 bps), had desirable on- and off-target scores^89^, and had no perfect match off-targets in the human genome (UCSC Blat)^90^. The ssODN repair templates were designed for each gRNA following optimal template design guidelines (Richardson et al.^91^). Specifically, ssODNs were ∼130 nucleotides in length with 36 bp overhang on the protospacer-adjacent motif (PAM)-distal side of the DNA cut site and a 91 bp overhang on the PAM-proximal side; the ssODNs were complementary to the first released DNA strand. The ssODNs (IDT Ultramer) were synthesized with 3 phosphorothioated bases on each end for all cell lines (except HEK293FT, where unmodified Ultramers were used) (Supplementary Table 4).

For TruEditor HDR experiments using the GFP to BFP reporter system, HEK293FT-GFP cells were electroporated using the SF Cell Line 4D-nucleofector kit (Lonza V4XC-2024) in 16-well format using HEK293FT electroporation program (CM-130). 5 × 10^5^ cells were nucleofected with 500 ng gRNA plasmid, 1 µg of lentiviral TruEditor or Cas9-only plasmid, and 0.35 µL of ssODN resuspended to 100 µM concentration (IDT Ultramer). 3-4 independent nucleofection reactions were performed for each construct, preparing fresh reagents each time. After 24 hours of recovery, cells were selected for 48-72 hours in 1 µg/mL puromycin. Following selection, cells were cultured for an additional 24 hours, then harvested and prepared for flow cytometry.

For TruEditor HDR rate quantification by targeted amplicon sequencing, HEK293FT, A549, and BT16 cells were nucleofected using the SF Cell Line 4D-nucleofector kit (Lonza V4XC-2024) with the CM-130 pulse program. For MDA-MB-231 cells, the SE Cell Line 4D-nucleofector kit (Lonza V4XC-1032) was used with the CH-125 pulse program. For HEK293 and A549 cell lines, 2.5 × 10^5^ cells were used per reaction. For MDA-MB-231cell lines, 5 × 10^5^ cells were used, and for BT16, 1 × 10^6^ cells were used. In every case, 400 ng of gRNA plasmid, 800 ng of CMV-driven TruEditor or Cas9-only plasmids, and 0.4 µL of ssODN resuspended to 100 uM concentration was used. 3-4 independent nucleofection reactions were performed for each construct, preparing fresh reagents each time. Cells were allowed 20 - 45 minutes recovery in fresh media after electroporation before culturing in 24-well plates with passaging for 5 days prior to gDNA extraction.

For prime editing experiments, 2.5 × 10^5^ HEK293FT cells were nucleofected with 800 ng of PEmax expression plasmid (Addgene 406249) and 200 ng engineered prime editing guide RNA (epegRNA) expression vector (Addgene 174039) using the SF Cell Line 4D-nucleofector kit (Lonza V4XC-2024) in 16-well format using HEK293FT electroporation program (CM-130). For flow cytometry, readouts were performed 6 days post-treatment. For amplicon sequencing, DNA was extracted at 4 and 8 days post-treatment.

For HUES66 cells, nucleofections were performed using the P3 primary cell line kit (Lonza V4XP-3024) using the CB-150 program. 2.5 × 10^5^ cells were mixed with 1 pmol Cas9 or TruEditor mRNA, 20 pmol synthetic gRNA, and 20 pmol end-modified ssODN per reaction. Cells were recovered in E8 media for 30 minutes at 37°C prior to plating in E8 with CEPT. Cells were cultured for at least 5 days before readout by flow cytometry.

For primary human CD8 T cells, the *CD19* CAR donor was generated by PCR of a *CD19* CAR construct using Q5 HiFi Polymerase (NEB M0491L). To amplify the double-stranded *CD19* donor, we performed 50 × 100 µL reactions with Q5 (30 s at 98°C, 30 cycles × [10 s at 98°C, 20 s at 98°C, 1 min at 72°C), 2 min at 72°C). The PCR product (5 mL) was pooled and then purified using AMPure XP beads (Beckman A63880) using 2 rounds of 0.5× left-sided SPRI to remove primers. Nucleofections were performed three days after T-cell isolation and activation using the P3 primary cell kit with the EH-115 program. The first SPRI was eluted in 500 µL and the second SPRI was eluted in 50 µL. In each case, 1 × 10^6^ T cells were treated with 2 pmol Cas9 or TruEditor mRNA, 40 pmol synthetic gRNA, and 3 µg of double-stranded *CD19* CAR donor template per reaction. After nucleofection, cells were recovered in T cell medium (TCM) which consisted of ImmunoCult T-cell Expansion Media (STEMCELL 10981) with 10 ng/mL human recombinant IL-2 (STEMCELL 78036). We added 150 µL of TCM to each well of the nucleofector strip and placed the cells in the incubator (37 °C, 5% CO_2_) for 30 minutes. After that, cells were transferred to 48-well plates in TCM media and cultured for at least 5 days before readout by flow cytometry.

### Flow cytometry and precise repair quantification for editing experiments

GFP and BFP positive cells were detected by flow cytometry (Sony SH800 or Miltenyi MACSQuant Analyzer 10). Cells were gated by forward and side scatter and signal intensity to remove cellular debris and multiplets. For each sample, at least 10,000 cells were analyzed for GFP and BFP signal. GFP and BFP gates were determined with reference to positive and negative controls for both markers, and precise editing was calculated as follows: the proportion of BFP-positive and GFP-negative cells after treatment divided by the total proportion of GFP-negative cells minus background (the proportion of GFP-negative cells in the no-transfection control).

TCR and CD19 CAR positive T cells were detected by flow cytometry (Miltenyi MACSQuant Analyzer 10). 2.5 × 10^5^ cells were stained with 1 µL APC-labeled anti-TCR alpha/beta mouse monoclonal antibody (BioLegend 306717), 1 µL PE-labeled anti-CD19 (FMC63 scFv) CAR mouse monoclonal antibody (ACROBiosystems FM3-PY54A2), and 0.04 µL of violet live/dead cell stain (Thermo L34963) in a total of 25 µL PBS for 30 minutes at room temperature. After washing 2x in PBS, cells were read out by flow cytometry and gated based on forward and side scatter to remove cellular debris and multiplets and live/dead positivity to isolate the living cell population. For each sample, at least 10,000 cells were analyzed for TCR and CD19 CAR signal. Gates were determined with reference to positive and negative controls for both markers, and precise editing was calculated as follows: the proportion of CD19 CAR-positive and TCR-negative cells after treatment divided by the total proportion of TCR-negative cells minus background (the proportion of TCR-negative cells in the no-transfection control).

### Targeted amplicon sequencing to quantify precise repair

Cells in 24-well plates were washed in PBS then lifted using 100 µL TrypLE (Thermo 12604039) before being transferred to 96-well PCR plates. Cells were centrifuged, and the supernatant was removed. 50 µL of QuickExtract (Lucigen QE09050) was added to each well and the plates were vortexed and incubated at 65 °C for 15 minutes, 68 °C for 15 minutes, and 98 °C for 5 minutes. The samples were stored at −20 °C until amplification.

A 2-step PCR protocol was used to amplify target loci from the gDNA extracts and attach barcodes, adapters, and stagger sequences for Illumina sequencing. For PCR 1, 500 ng of DNA was used as the template for a 25 µL Q5 PCR reaction with 20 cycles. PCR 1 products were run on a 2% 96-well E-Gel (Thermo G402022) and normalization was performed based on band intensity (BioRad Image Lab v5.2.1). Depending on the intensity of the PCR 1 band, 1 - 4 µL of PCR 1 product was used as input for PCR 2 with 8 additional cycles. PCR2 products were again run on 2% 96-well E-Gels, normalized, and pooled. The resultant amplicon pool was double-sided SPRI purified (Beckman-Coulter AMPure) at 0.5× and 0.7×. The final purified library was sequenced on a MiSeq (Illumina).

Sequencing reads were demultiplexed using Illumina i7 barcodes on the PCR2 reverse primers with bcl2fastq (v2.20.0.422), and then by in-read barcodes. Reads were trimmed on both sides based on PCR2 primer-binding sequences and quality filtering was performed simultaneously with cutadapt (v4.0)^92^. Additional quality control steps were taken prior to HDR quantification including fastqc (v0.11.3)^93^ and multiqc (v1.8)^94^ to ensure read quality, and genome alignment with bowtie2 (v2.5.1)^95^ to ensure amplicons mapped to the intended target loci. Next, CRISPResso2 (v2.2.12)^96^ was used in batch mode to quantify precise HDR and indel rates with default parameters plus the following: `-min_bp_quality_or_N 30 -w 5`. HDR rates were quantified as the number of precise HDR reads over the total number of edited reads (precise HDR, imprecise HDR, and all indels).

### Immunofluorescence

For DNA damage immunofluorescence assays, HEK293FT and A549 cells stably expressing Cas9 or TruEditors were seeded on clear-bottom, black 96-well plates (Corning 3904) at a density of 4 × 10^4^ and 5 × 10^4^ cells/well, respectively. For HEK293FT, plates were coated prior to cell seeding with 0.1 mg/mL poly-D-lysine (Sigma-Aldrich P1024) and washed. Cells were treated with doxorubicin (0.5 μM, MedChem Express 23214-92-8), or DMSO (Sigma-Aldrich 67-68-5) for 4 hours to induce DNA damage. Cells were fixed in 4% paraformaldehyde (Electron Microscopy Sciences 15710-S) or washed with media and fixed after 24 hours of recovery. DNA-PKci treatments NU7441 (4 μM, Selleck Chemicals 503468-95-9) and M3814 (2 μM, Selleck Chemicals 1637542-33-6) were added to cells during drug treatment and recovery.

The plates were washed with PBS and cells were permeabilized with Triton X-100 (0.1%). Next, plates were blocked (3% BSA and 0.1% Tween-20 in PBS) for 1 hour then incubated with mouse anti-yH2A.X (1:1000, EMD Millipore 05-636) and rabbit anti-53BP1 (1:1000, Novus Biologicals NB100-304) at 4°C overnight. Next, the plates were washed 3 times with PBS and incubated with Alexa Fluor Plus 488-conjugated goat anti-mouse IgG secondary antibody (1:500, Invitrogen A32723), Alexa Fluor 555-conjugated goat anti-rabbit IgG secondary (1:500, Invitrogen A-21428), and Hoechst 33342 (1:2000, Thermo H3570) at room temperature for 1 hour and then washed in PBS 3 times.

For the DNA damage assays, immunofluorescence images were acquired using a widefield fluorescence microscope (Keyence BZ-X) and analyzed using CellProfiler (v4.2.6)^97^ to quantify nuclear fluorescence and foci. To ensure data quality, each Hoechst image was first evaluated for sharpness using the `MeasureImageQuality` module, with a blur metric computed at a spatial scale of 20 pixels for HEK293 and 25 pixels for A549 cells. Images below the quality threshold were excluded from downstream analysis. To correct for non-uniform illumination across wells and fields, we applied the `CorrectIlluminationCalculate` module using a block size of 20 pixels and a polynomial fitting smoothing method to generate illumination correction functions. These were applied to each image using the `CorrectIlluminationApply` module via subtraction. Nuclei were identified in the Hoechst channel using the `IdentifyPrimaryObjects` module with a pixel diameter range of 20–100, threshold scale of 0.001–0.01, and a 2-class thresholding strategy.

To enhance detection of DNA damage foci within nuclei, the yH2AX and 53BP1 channels were processed with the `EnhanceOrSuppressFeatures` module using speckle enhancement at a feature size of 10 pixels, followed by masking of extranuclear signal with the `MaskImage` module. Foci were then segmented using `IdentifyPrimaryObjects` with a pixel diameter of 1–14, threshold scale range of 0.0001–0.5, and 2-class thresholding. These objects were related to their parent nuclei using the `RelateObjects` module. Fluorescence intensity for Hoechst, yH2AX, and 53BP1 channels was quantified within each nucleus using the `MeasureObjectIntensity` module. Additionally, the number, integrated intensity, and size of foci were quantified for each nuclear object in both the yH2AX and 53BP1 channels. For visualization, representative images were processed in Fiji (ImageJ2; https://imagej.net/software/fiji/)^98^ with uniform adjustments to brightness, contrast, and gamma applied consistently within each imaging channel.

For percent recovery of 53BP1 foci after treatment with 0.5 µM doxorubicin, we first computed the average foci count per nuclei for each biological replicate of each treatment (vehicle, NU7441, M3814, Cas9 or TruEditor transduction) — for both the 4-hour doxorubicin and 4-hour doxorubicin plus 24-hour recovery conditions (median of 612 nuclei for A549 and 520 nuclei for HEK293FT per replicate). Next, the average 53BP1 foci count per nuclei in the 24-hour recovery condition for each replicate was subtracted from the treatment-matched 4-hour (no recovery) condition. For this step, we used the average (over biological replicates) for the treatment-matched 4-hour (no recovery) conditions; we term this quantity the *TMA_Doxo4hr*. These differences (*TMA_Doxo4hr* minus *4-hour doxorubicin, 24-hour recovery*) were normalized by the *TMA_Doxo4hr* (and multiplied by 100) to compute a percent change. For percent recovery of γH2AX, we performed the same calculations as above but using integrated γH2AX intensity per nuclei instead of 53BP1 foci.

For FLAG affinity tag immunofluorescence, HEK293FT cells were seeded 7 days post-nucleofection on poly-D-lysine-coated 96 well plates at 1 × 10^5^ cells/well. Cells were grown for 36 hours, fixed, permeabilized, and blocked according to the above procedure. The plates were incubated with mouse anti-FLAG (1:500, Sigma-Aldrich F1804) and rabbit anti-PARP1 (1:400, Thermo 22999-1-AP) at 4 °C overnight. Next, the plates were washed and incubated with Alexa Fluor Plus 488-conjugated goat anti-mouse IgG secondary antibody (1:500), Alexa Fluor 647-conjugated donkey anti-rabbit IgG secondary antibody (1:500, Thermo A31573) and Hoechst 33342 (1:2000) at room temperature for 1 hour and then washed in PBS 3 times.

For stem cell GFP/BFP imaging and Oct3/4 immunofluorescence, edited HUES66-oNGN-GFP hESCs were seeded at 15 × 10^3^ cells per well in 100 µL E8 medium (Thermo A1517001) supplemented with CEPT cocktail (Tocris 7991) in Geltrex-coated 96-well plates (Corning 3904). The following day, medium was replaced with E8 medium. After 24 hours, cells were fixed in 4% paraformaldehyde (Electron Microscopy Sciences 15710-S) for 15 minutes at room temperature and washed 3 times with PBS. Cells were permeabilized with 0.1% Triton X-100 in PBS for 10 minutes, washed twice with PBS, and blocked with 3% BSA and 0.1% Tween-20 in PBS (blocking buffer) for 1 hour at 4 °C. Primary antibody staining was performed using anti-Oct3/4 (1:500, Santa Cruz Biotechnology sc-5279) diluted in blocking buffer and incubated overnight at 4 °C. The next day, cells were washed 3 times with PBS and incubated with Alexa Fluor 647-conjugated goat anti-mouse IgG secondary antibody (1:800, Thermo A-21235) for 1 hour at room temperature. Following 3× PBS washes, fluorescent images (GFP, BFP, and Oct3/4) were acquired (Keyence BZ-X). Nuclei were then counterstained with Hoechst 33342 (1:2000, Thermo H3570) diluted in blocking buffer for 15 minutes at 4 °C, washed twice with PBS, and re-imaged at the same positions.

### Immunoprecipitation and western blots

5 × 10^5^ HEK293FT cells were plated into 6-well plates and allowed to adhere overnight. On the following day, 2 µg of FLAG-tagged TruEditor or Cas9-only expression vectors was transfected using Lipofectamine 3000 (Thermo L3000008). Protein pellets were harvested at 24 hours after transfection as follows: HEK293 cells were lifted with 250 µL TrypLE (Thermo 12604039), quenched with 1 mL D10 media, and pelleted at 500 × g for 3 minutes. Pellets were combined with 170 µL pre-cooled immunoprecipitation lysis buffer (Thermo 87787) supplemented with 1% protease inhibitor cocktail (Selleck Chemicals B14001), mixed vigorously, and incubated for 30 minutes on ice with frequent inverting. Next, the lysate solutions were pelleted at 10,000 x g for 10 minutes and 50 µL of the supernatant was set aside as an input control; the remaining supernatant was transferred into a fresh tube containing 10 µL of pre-washed anti-FLAG M2 magnetic beads (Millipore M8823) according to manufacturer’s protocol. Samples were incubated with anti-FLAG beads overnight at 4 °C in a roller-shaker. Finally, samples were washed according to manufacturer’s protocol and samples were analyzed by mass spectrometry or western blotting, as described below.

For protein blots, samples were denatured in NuPAGE LDS Sample Buffer (Thermo P0007) supplemented with 100 mM DTT (Cayman 700416) for 10 minutes at 70 °C. Denatured samples and PageRuler prestained protein ladder (Thermo 26616) were separated in Novex 4-12% Tris-Glycine mini gels (Thermo XP04125) in 1× MOPS running buffer (Thermo NP0001) for 75 minutes at 160 V, then transferred to nitrocellulose membranes (BioRad 1620112) in 1× Tris-Glycine transfer buffer (Thermo LC3675) supplemented with 10% methanol for 1 hour at 33 V.

Membranes were blocked in 5% milk (Research Products International M17200) dissolved in 1× TBS with 0.1% Tween-20 (TBS-T) for 60 minutes. The membranes were then incubated overnight at 4 °C with one of the following primary antibodies: mouse anti-*Sp*Cas9 (Cell Signaling 14697, 1:2000), rabbit anti-TOP3A (Proteintech 14525-1-AP, 1:2000), rabbit anti-RMI1 (Proteintech 14630-1-AP, 1:500), or rabbit anti-ACTB (Proteintech 81115-1-RR) in 5% BSA (VWR AAJ65097) dissolved in TBS-T. The blots were washed and then incubated with IRDye 800CW donkey anti-rabbit or anti-mouse secondary antibodies (LI-COR 925-32213, 925-32212, 1:10,000 dilution in 5% BSA/TBS-T) for 60 minutes at room temperature, washed 3 times with TBS-T and visualized using an Odyssey CLx imaging system (LI-COR).

### Mass spectrometry for affinity proteomics

MS sample processing and basic enrichment analysis were performed by the NYU Langone Proteomics Laboratory (SCR_017926). Bead-bound protein samples were washed 3 times with 100 mM NH_4_HCO_3_ (pH 8.0) and reduced with 2.5 μL 0.2 M dithiothreitol at 57 °C for 1hour. Next, the samples were alkylated with 2.5 μL 0.5M Iodoacetamide (IAM) for 45 minutes at room temperature in the dark and digested overnight on a shaker at room temperature with 500 ng of sequencing-grade modified trypsin (Promega). Following digestion, peptide samples were separated from magnetic beads using a magnetic rack. Samples were acidified to a final concentration pH of 2.0 using 10% trifluoroacetic acid (TFA).

One-third of each sample was analyzed individually by liquid chromatography–tandem mass spectrometry (LC–MS/MS). Samples were separated online using an Evosep One (EVOSEP 1112 performance column 15 cm x 75 µm packed with 1.9 µm ReproSil-Pur C18 beads). Peptides were gradient eluted from the column directly to the Thermo Scientific Orbitrap Eclipse Tribrid mass spectrometer using an 88 min gradient (Thermo), in which solvent A was 2% acetonitrile/0.5% acetic acid and solvent B was 80% acetonitrile/0.5% acetic acid. High resolution full MS spectra were acquired with a resolution of 120,000, an automatic gain control (AGC) target of 4 × 10^6^, with a maximum ion time of 50 ms, and scan range of 400 to 1500 m/z. All MS/MS spectra were collected using the following instrument parameters: in the ion trap, AGC target of 2 × 10^4^, maximum ion time of 22 ms, one microscan, 1.5 m/z isolation window, fixed first mass of 110 m/z, and normalized collisional energy (NCE) of 30. MS/MS spectra were searched against a Uniprot human database using Sequest within Proteome Discoverer 2.5. SAINT analysis^99^ was performed on the resulting peptide spectral match (PSM) data to identify bona fide interaction partners using the Cas9 pulldown as a control.

Raw PSM values were used to compute cross-sample Pearson correlations. Hierarchical clustering and heatmap visualization were performed on the correlation matrix using the pheatmap package. For the remainder of analyses, bona fide interaction partners for TruEditor proteins were defined as those having an average PSM greater than 2.5, fold-change over Cas9 above 10, and SAINT FDR-adjusted *P* < 0.05. Gene Ontology (GO) FDR-adjusted *P* values were computed for the entire dataset of enriched proteins. Protein-protein interaction diagrams were generated with Cytoscape (v3.10.3)^100^ and GO enrichment was performed using the stringApp (v2.2.0)^101^.

### In vitro mRNA transcription

Polyadenylated Cas9 or TruEditor mRNA containing fully substituted N1-methyl pseudouridine (BOC Sciences B2706-358101) and a Cap 1 Structure (m7G(5′)ppp(5′)(2′OMeA)pG, BOC Sciences B1370-351546) was synthesized through *in vitro* transcription (IVT) using a PCR amplicon for templated synthesis. First, the IVT template was generated by amplifying constructs cloned into pT7 vectors using 25 cycles with Q5 high-fidelity polymerase (NEB M0491L) with a forward primer correcting the mutated T7 promoter sequence and a reverse primer with a poly-thymine (T) handle to generate poly-adenylated mRNA. The amplified sequence spans the T7 promoter to the 3’UTR. IVT templates were concentrated and purified using AMPure XP beads (Beckman A63880) with a 0.5X SPRI cleanup.

Next, reaction mixes containing 5 mM each of pseudouridine and NTPs (adenosine-, cytidine-, guanosine-5′-triphosphate, NEB E2040S), 4 mM mRNA Cap AG (Cap1), 2.5 µg of the purified template, 1 unit/µL murine RNase inhibitor (NEB M0314S), 0.002 units/µL inorganic yeast pyrophosphatase (NEB M2403S), and 50 units/µL T7 RNA polymerase (NEB M0251L) were assembled in 1x transcription buffer (100 µL total volume) and incubated for 3 hours at 37 °C. For in vitro transcription, we first made a 10**×** transcription buffer: 400 mM Tris-HCL (IBI Scientific IB70162), 100 mM DTT (Sigma 43816), 20 mM spermidine (Sigma 85558), 0.02% Triton X-100 (Sigma 93443), and 165 mM magnesium acetate (Sigma 63052). Next, DNase treatment was performed after diluting the IVT reaction volume to 480 µL in molecular biology grade water (Cytiva SH30538) and adding 50 units of DNase I (NEB M0303S), 4.8 µL of 200 mM CaCl_2_ (Sigma 21115), and incubating for an additional 20 minutes at 37 °C. Finally, mRNA was purified using a Monarch mRNA purification kit (NEB T2050L). The ssRNA concentration was determined by nanodrop and all aliquots were stored at −80 °C prior to use in editing experiments.

### Primary human T cell isolation and culture

PBMCs were isolated from healthy donor Human Peripheral Blood Leukapheresis Packs (Leukopak, Stemcell 200-0092). Leukopaks were collected in 50 mL tubes, washed and resuspended in RPMI media with L-glutamine (Cytiva SH30255). Red cell lysis was performed using 25 mL of a 1**×** red cell lysis buffer solution (Tonbo Bioscience TNB-4300-L100) for 10 minutes at 37 °C. Following lysis, cells were washed and resuspended in BamBanker Freezing Medium (Bulldog Bio BB01) at a concentration of 100 **×** 10^6^ cells / mL. Cells were first transferred to −80 °C for 24 hours before moving to liquid nitrogen for long term storage.

Cryopreserved PBMCs were defrosted in a 37 °C water bath for ∼2 minutes before being transferred dropwise into RPMI media supplemented with 10% FetalClone III serum by volume (R10 media) and 10 ng/mL Human Recombinant IL-2 (Stemcell 78036.3). The cells were washed by centrifugation and resuspension and passed through a pre-wetted 40 µm cell strainer (Corning 352340) into a fresh 50 mL tube. Next, CD8+ T cells were isolated from PBMCs using the EasySep Human CD8 Positive Selection Kit II (Stemcell 17853) and plated in 24-well plates at a concentration of 1 **×** 10^6^ cells / mL in T cell medium (TCM) which consisted of ImmunoCult T-cell Expansion Media (STEMCELL 10981) with 10 ng/mL human recombinant IL-2 (STEMCELL 78036); they were immediately activated with 25 µL per well Human CD3/CD28 T cell Activator (Stemcell 10991). CD8+ T cells were maintained in TCM at 37 °C and 5% CO_2_ with regular passaging.

### CAR T tumor killing assay

CD19+ Nalm6 cells expressing EGFPd2PEST-NLS^41^ were cultured in R10 with 2 µg/mL puromycin, which was removed from the medium 24 hours prior to CD8+ T cell co-culture. 2.5 **×** 10^4^ Nalm6-GFP cells per well were plated in a black-walled 96-well plate (Corning 3904) in TCM without IL-2. Next, 1.25 **×** 10^4^ T cells 2 weeks post-nucleofection were added to the wells and mixed thoroughly by pipetting and the mixed co-culture was topped up to a total volume of 150 µL per well using TCM without IL-2. Live-cell imaging was performed at 20**×** magnification acquiring 4 images per well every 2 hours for 120 hours using Incucyte (Sartorius). For each well, the integrated GFP intensity was normalized to the 2-hour time point.

### Protein structure prediction using AlphaFold

Wild-type amino-acid sequences were obtained from UniProt or from sequences of cloned TruEditor vectors. Structural predictions and interaction models were generated using the AlphaFold Server (AlphaFold3). For each query, only the top-ranked prediction (based on AlphaFold’s internal ranking score) was used. Model confidence was assessed using the predicted template modeling score (pTM) for global structure and the interface pTM (ipTM) for inter-chain interactions. 3D structural renderings were visualized using Mol* Viewer^102^.

### Data representation and statistical analysis

All bar plots represent the mean of at least 3 biological replicates (independent nucleofection reactions from individually harvested cell populations) with individual replicates plotted as points and error bars representing s.e.m. In all cases when comparing TruEditors to a control group, ANOVA was performed with post-hoc pairwise comparisons conducted using Tukey’s HSD test. Statistical analyses were performed in R (v4.4.1). Flow cytometry data were analyzed using FlowJo (v10.10.0).

## Acknowledgments

We thank the entire Sanjana laboratory for their support and advice. We also thank NYU Biology Genomics Core for sequencing resources and acknowledge the Zegar Family Foundation for their generous support of the Core. We also thank the Proteomics Laboratory (SCR_017926), which is subsidized in part by NYU Langone and the Laura and Isaac Perlmutter Cancer Center support grant P30CA016087 from the National Cancer Institute.

## Author contributions

W.H.B., Z.D., A.S. and N.E.S. designed the study. Z.D. and L.L. performed the CRISPR screen. W.H.B., S.J.M. and N.E.S. performed CRISPR screen analyses. W.H.B. designed and tested TruEditors. W.H.B., C.Y.C. and G.I.C. cloned plasmids. W.H.B., R.E.Y., A.H.Y.S. and Z.D. designed donor templates. S.J.M., P.D.D. and W.H.B. performed protein structure visualizations. W.H.B. and C.J. generated TruEditor mRNA. W.H.B. and P.D.D. performed DNA repair experiments. W.H.B., A.R. and L.K. performed stem cell experiments. W.H.B., S.J.M. and M.D. performed western blotting and affinity proteomics. W.H.B., S.J.M. and N.E.S. wrote the manuscript with input from all authors. N.E.S. supervised the study.

## Materials availability

Plasmids will be made available on Addgene.

