## Supplementary Figures for "Rational design of synthetic proteins using a genome-scale CRISPR screen"

### Supplementary Tables

**Supplementary Table 1.** Fold-change of gRNAs in the CRISPRa screen for sorted HDR and NHEJ populations compared to pre-sort cells.

**Supplementary Table 2.** Gene-level analysis of the CRISPRa screen using Robust Rank Aggregation.

**Supplementary Table 3.** TruEditor amino acid sequences and mRNA sequences.

**Supplementary Table 4.** Precise editing gRNA sequences, ssODN sequences, and primer sequences.

**Supplementary Table 5.** TruEditor affinity proteomics by construct.

### Supplementary Movies

**Supplementary Movie 1.** AlphaFold3 predicted co-folding of the Cas9-XTEN-BARD1 TruEditor with AP-MS identified BRCA1 binding partner.

**Supplementary Movie 2.** AlphaFold3 predicted co-folding of the Cas9-XTEN-FAN1 TruEditor with AP-MS identified PMS2 and MLH1 binding partners.

**Supplementary Movie 3.** AlphaFold3 predicted co-folding of the Cas9-XTEN-ZRANB3 TruEditor with AP-MS identified PCNA binding partner.

**Supplementary Movie 4.** AlphaFold3 predicted co-folding of the Cas9-XTEN-ZNF146 TruEditor with AP-MS identified YY1 binding partner.

**Supplementary Movie 5.** AlphaFold3 predicted co-folding of the Cas9-XTEN-$BLM_{2\text{-}194}$ TruEditor with AP-MS identified RMI1 and TOP3A binding partners.

### Supplementary Figures

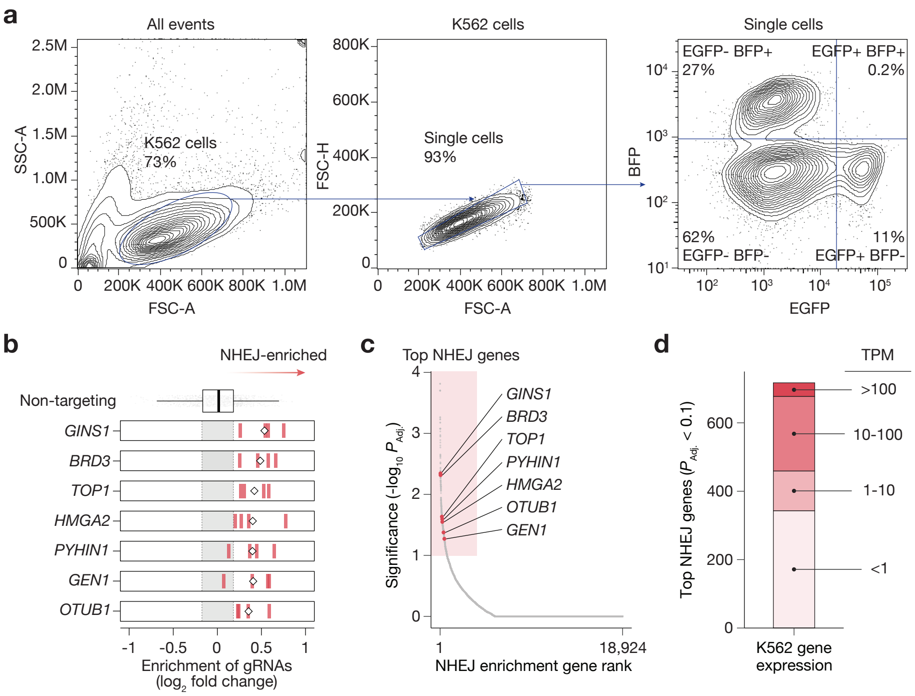


**Supplementary Figure 1. The genome-scale CRISPR activation (CRISPRa) screen also identifies modulators of non-homologous end-joining (NHEJ). a**, Flow cytometry gating for genome-scale CRISPRa screens in human K562-EGFP cells. **b**, Normalized enrichment in the NHEJ population for the four most enriched CRISPRa gRNAs targeting the indicated genes. The fold-change of each gRNAs is shown as blue lines and the average is indicated by the diamond. **d**, Gene ranks and significance for NHEJ enrichment with robust rank aggregation (RRA) using the four most enriched gRNAs. The highlighted region indicates genes with FDR-adjusted RRA $P_{adj} < 0.1$. **e**, RNA expression in K562 cells of the top enriched NHEJ genes (FDR $P_{adj} < 0.1$).

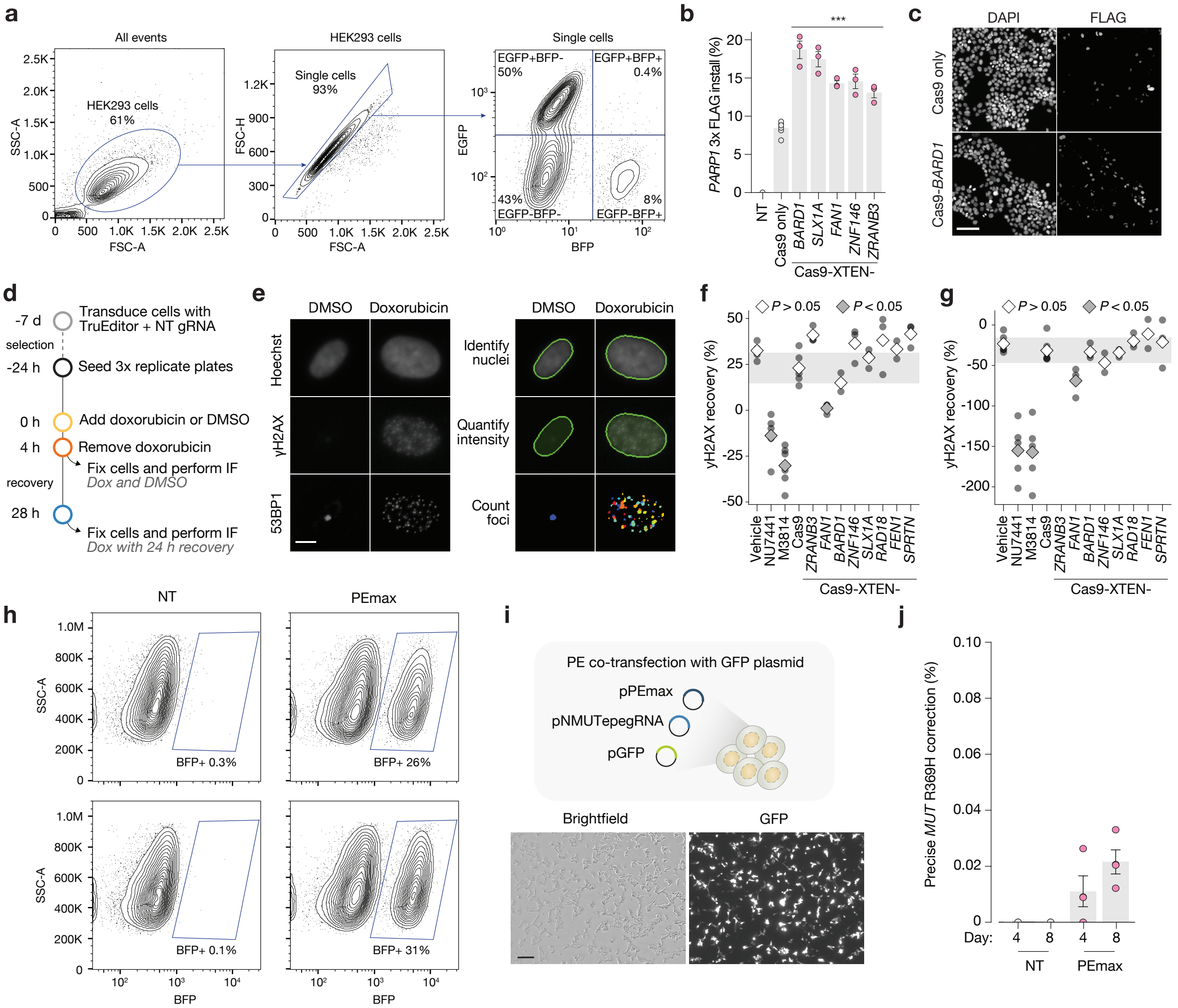


**Supplementary Figure 2. Most TruEditors do not change global DNA damage responses and prime editing at the *MUT* locus results in near-zero editing. a**, Flow cytometry gating for GFP-to-BFP precise editing assay in human HEK293-EGFP cells. **b**, Precise knock-in of 3xFLAG affinity tag at the N-terminus of the *PARP1* gene in HEK293 cells with Cas9 only and various TruEditors. **c**, Representative anti-FLAG immunofluorescence (IF) and nuclear (DAPI) stain of human HEK293 cells at the *PARP1* locus using Cas9 or a Cas9-BARD1 TruEditor. Scale bar, 50 μm. **d**, Schematic timeline for DNA damage and recovery assay to interrogate effects of TruEditor expression on global DNA repair. **e**, Representative IF of human lung A549 cells treated with vehicle (DMSO) or doxorubicin for markers of DNA damage (yH2AX and 53BP1) (*left*) and with image processing overlay (*right*). Scale bar, 10 μm. **f**, Recovery of yH2AX foci in human lung A549 cell transduced with TruEditors after doxorubicin removal. We included non-transduced cells treated with vehicle/DMSO (negative control) or DNA-PK inhibitors NU7441 or M3814 (positive control). **g**, Same quantification as in panel *f* in human HEK293 cells. **h**, Precise GFP to BFP conversion rates in human HEK293-EGFP cells using prime editing. **i**, Prime editing transfection by co-transfection of prime editing reagents and a GFP expression vector in HEK293 cells. Scale bar, 200 μm. **j**, Precise editing rates for correcting the *MUT* R369H mutation in human HEK293 cells using prime editing. For all comparisons, experiments had at least 2 biological replicates and significance was determined using a one-way ANOVA. Post-hoc pairwise comparisons were conducted using Tukey's HSD test (*$P < 0.05$, **$P < 0.01$, ***$P < 0.001$).

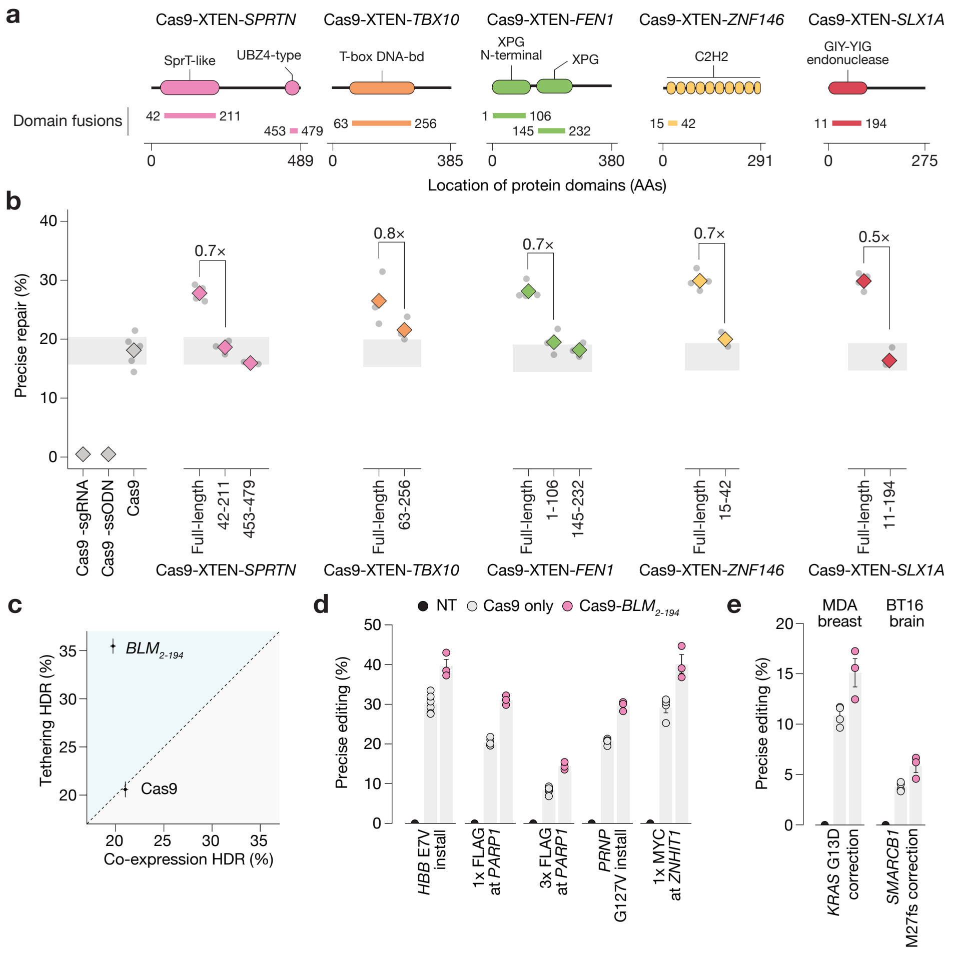


**Supplementary Figure 3. Additional minimal TruEditors tested using core protein domain fusion. a**, Linear representation of annotated protein domains from the indicated proteins. Cloned domains or combinations of domains are indicated as horizontal lines and their lengths given in amino acids. **b**, Precise repair (GFP-to-BFP editing) rates for the indicated minimal or full-length TruEditors. TruEditor constructs were nucleofected into human kidney HEK293-EGFP cells with a GFP-targeting gRNA and a single-stranded repair template encoding BFP-specific edits. Points represent three independent transfection replicates, and the diamond indicates the mean. Grey rectangle represents the 95% confidence interval for the Cas9 control. **c**, Comparison of $BLM_{2-194}$ TruEditor as direct Cas9 fusion versus co-expression (XTEN linker or 2A cleavage site). Points represent the mean of three replicates with error bars indicated SEM. **d**, Precise editing (amplicon sequencing) at the indicated locus with Cas9 only or a minimal $BLM_{2-194}$ TruEditor in human HEK293 cells. **e**, Precise editing (amplicon sequencing) at the indicated locus with Cas9 only or a minimal $BLM_{2-194}$ TruEditor in human breast cancer MDA-MB-231 or brain atypical teratoid/rhabdoid tumor (ATRT) BT16 cells. All experiments had at least 3 biological replicates.

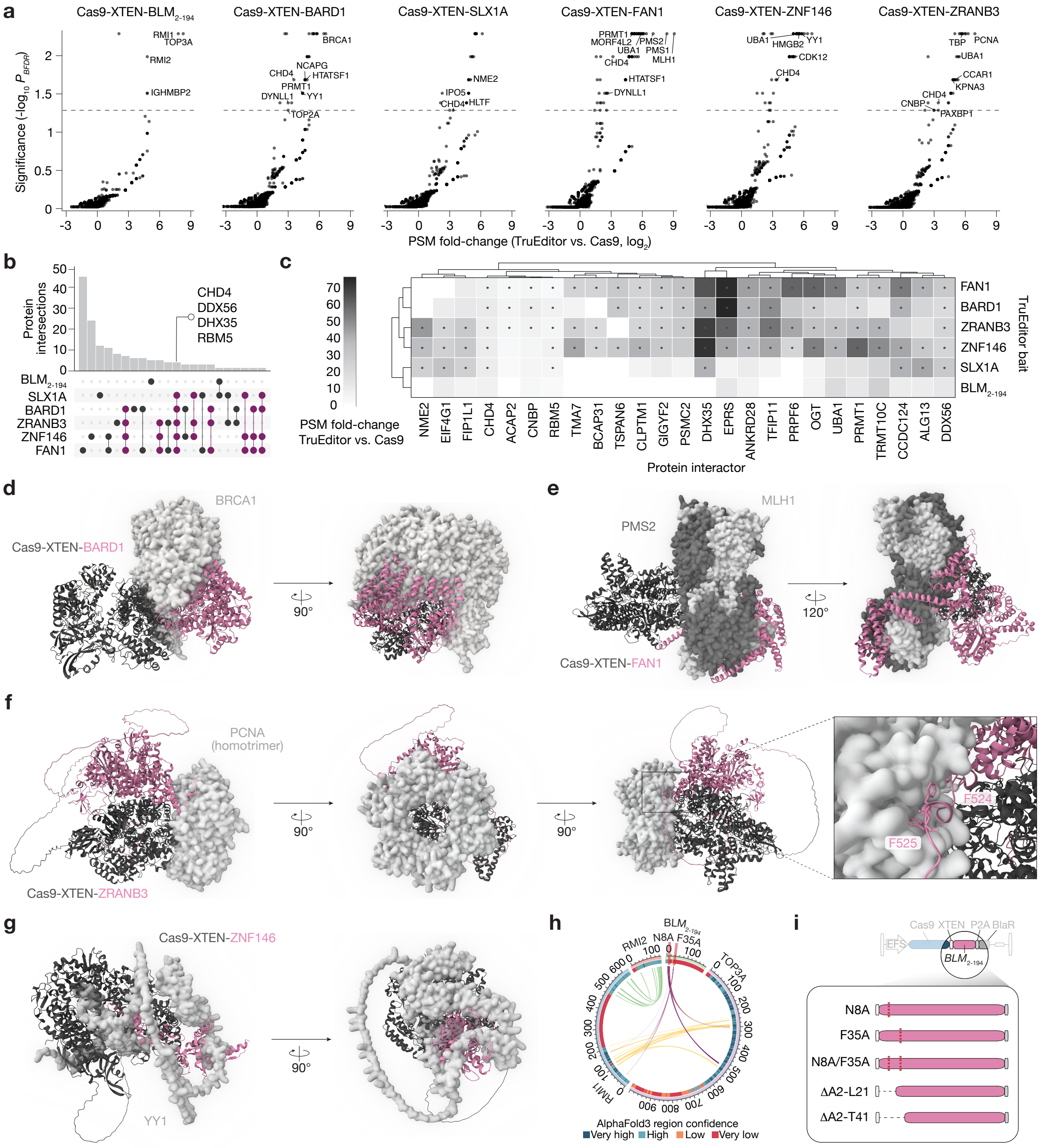

**Supplementary Figure 4. TruEditors coordinate distinct endogenous DNA repair proteins to improve precise gene editing. a**, Enrichment of proteins co-immunoprecipitated with the indicated TruEditor versus Cas9 only (anti-FLAG affinity proteomics-mass spectrometry [AP-MS]). Significance was determined by SAINT analysis[23] and the dotted line indicates Bayesian false-discovery rate (BFDR) $P_{BFDR} < 0.05$ (n = 2 - 3 biological replicates per TruEditor or Cas9). **b**, Overlap of AP-MS enriched proteins for the indicated TruEditors, where enrichment was determined against Cas9 only ($P_{adj} < 0.05$). **c**, Peptide Spectral Match (PSM) fold-change over Cas9 only for proteins enriched in 3 or more AP-MS datasets. Significant fold-change represented by black dots ($P_{adj} < 0.05$). **d-g**, AlphaFold3 predicted co-folding of the indicator

TruEditor with AP-MS identified binding partners. **h**, AlphaBridge[24] ribbon plot illustrating predicting interaction interfaces between the Cas9-*BLM*$_{2\text{-}194}$ TruEditor co-folded with complex partners *TOP3A*, *RMI1,* and *RMI2* using AlphaFold3. **i**, Schematic of Cas9-*BLM*$_{2\text{-}194}$ mutants to ablate protein interactions identified by AP-MS.

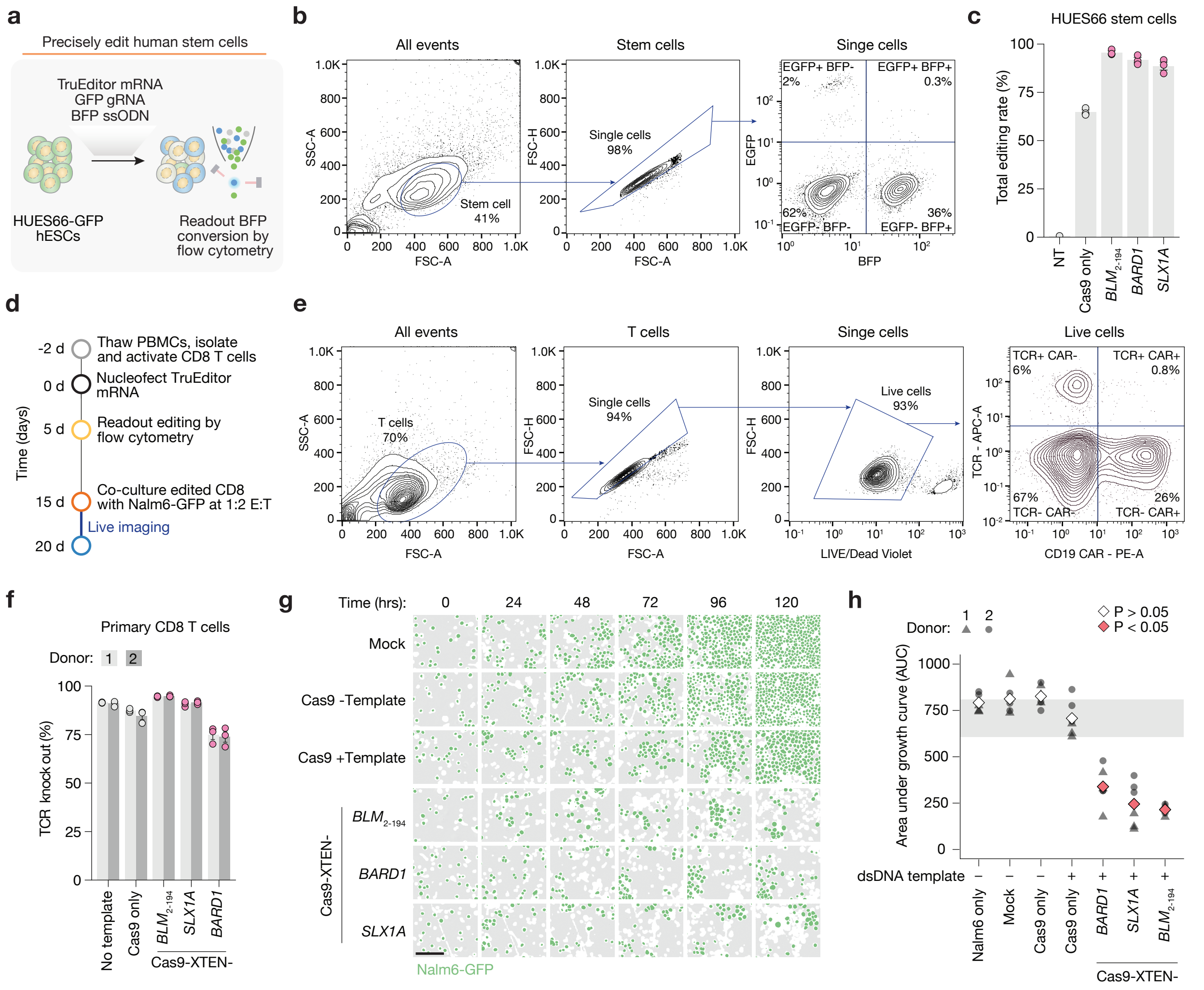

**Supplementary Figure 5. TruEditor messenger RNA (mRNA) delivery improves precise gene editing. a**, HUES66-EGFP cells were nucleofected with Cas9 or TruEditor mRNA, a GFP-targeting synthetic gRNA, and a BFP ssODN repair template. Following 5 days in culture, precise editing (GFP-to-BFP conversion) was quantified by flow cytometry. **b**, Flow cytometry gating for GFP-to-BFP precise editing assay in HUES66-EGFP cells. **c**, Total editing rates in HUES66-EGFP cells nucleofected with TruEditor mRNA, GFP-targeting gRNA, and BFP ssODN ($n$ = 3 nucleofection replicates). **d**, Primary human CAR T cell engineering experimental timeline. **e**, Flow cytometry gating to quantify *CD19* CAR knock-in in primary human CD8 T cells. **f**, TCR knockout (total editing) rates in primary human T cells nucleofected with TruEditor mRNA, *TRAC*-targeting gRNA, and a 2.8 kb *CD19* CAR donor template ($n$ = 2 human donors with 3 nucleofection replicates each) . **g**, Representative images from edited primary CD8+ human T cells in co-culture with CD19+ Nalm6-GFP leukemia cells (effector:target [E:T] ratio of 1:2). Scale bar, 100 µm. **h**, GFP area under the curve (AUC) for co-culture (T cells with Nalm6-GFP) time course experiments ($n$ = 24 images per condition with 4 images per nucleofection and 3 nucleofections for each of 2 human primary T cell donors). Statistical significance was determined using a one-way ANOVA with Tukey's HSD post hoc test (*$P$ < 0.05, **$P$ < 0.01, ***$P$ < 0.001).
